# Safety learning induces postsynaptic potentiation of direct pathway spiny projection neurons in the tail of the striatum

**DOI:** 10.1101/2025.04.29.651089

**Authors:** Adrien Stanley, Se Joon Choi, Anika Frank, Emily Makowicz, Stefano Cataldi, N Shashaank, Eugene V. Mosharov, Clay Lacefield, Maria Concetta Miniaci, David Sulzer

## Abstract

The association of a sensory cue with an outcome is a crucial step in learning to identify safe versus threatening situations. Here we assessed how learned sound-safety association alters neuronal activity in tail of the striatum (TS), where auditory cortical and thalamic inputs converge. Prior to training, foot shock elicited responses by TS direct and indirect pathway spiny projection neurons (SPNs), while sound tones produced no response. However, once the sound association was learned, sound tones strongly activated TS SPNs, even when the animal was under anesthesia. This conditioned auditory response occurred concurrently with alterations of direct pathway SPNs in the TS, including increased responses to cortical and auditory thalamic inputs, increased excitatory response with an enhanced ratio of NMDA to AMPA receptors, decreased responses to inhibitory input, and increased dendritic spines. This convergence of postsynaptic changes provides responses to relevant auditory cues during associative learning.

**Teaser:** *Sound-safety association enhances response to learned sound in striatal neurons via postsynaptic mechanisms*

## INTRODUCTION

Most chordate species use sound for communication (*1*) and the ability to learn the association of sound with experience is central to many behaviors. For example, learned calls by prairie dogs are required for the colony to safely forage and avoid danger (*2*). Anxiety and post-traumatic stress disorder (PTSD) can arise from aberrations in sound-safety and sound-threat associations (*3–7*). Here we analyze mechanisms underlying sound-safety association. We use a safety conditioning paradigm in which a conditioned stimulus (CS), an auditory cue, becomes capable of suppressing responses elicited by threatening cues (i.e., freezing behavior) (*3, 11–14*).

The striatum plays a central role in integrating sensory inputs from cortical and thalamic afferents to determine the appropriate behavioral output. The primary neurons of the striatum, known as spiny projection neurons (SPNs), send projections to downstream basal ganglia nuclei that control the sequencing of cortical motor signals transmitted to the spinal cord (*15*). A striatal subregion seemingly positioned to control learned sound association is the tail of the striatum (TS: also known as the auditory striatum), as TS activity is required for the acquisition and expression of auditory threat conditioning (*16, 17*) consistent with reports that TS local field potentials are enhanced following sound-safety association (*13*). Both auditory cortical and auditory thalamic (medial geniculate body) nuclei extend excitatory axonal projections to the TS (*18*). The auditory thalamic projections to the TS form synapses onto SPN dendrites (*19, 20*) where they modulate the response amplitude (*21*).

TS SPNs are comprised of “direct pathway” SPNs (dSPNs) that project to the substantia nigra reticulata (SNr) and globus pallidus internal (GPi), and “indirect pathway” SPNs (iSPNs) that project to the SNr/GPi via the globus pallidus externa (*22–25*). It was recently demonstrated that TS SPN activity is altered following sound-threat association (*16, 17*), with ventral TS dSPNs exhibiting increased response to threat-associated sounds, while iSPNs activity displayed relatively little change (*17*). However, the roles and synaptic modifications that might allow SPNs to encode sound-safety association are unknown.

## RESULTS

### Prior to training, foot shock but not sound cue activates TS SPNs

We first analyzed the responses of TS SPNs to foot shock and tone in naive mice. We injected AAV9-flex-GCaMP6f virus into the TS of D1:cre mice to measure responses by dSPNs (*26, 27*) and into the TS of A2A:cre mice to measure responses by iSPNs (*28, 29*).

After three weeks of viral expression, we transiently placed the mice in a new chamber (the “context”) and recorded GCaMP signals from optical fibers within the TS while playing a tone (1000 Hz square wave, 20 sec, 70 dB spl) or administering foot shocks (2 sec, 0.6 mA) (Figure 1A-C). In naive mice, the tones produced no detectable responses, while foot shock produced calcium responses by both TS dSPNs and iSPNs (Figure 1D-G).

**Figure 1.**
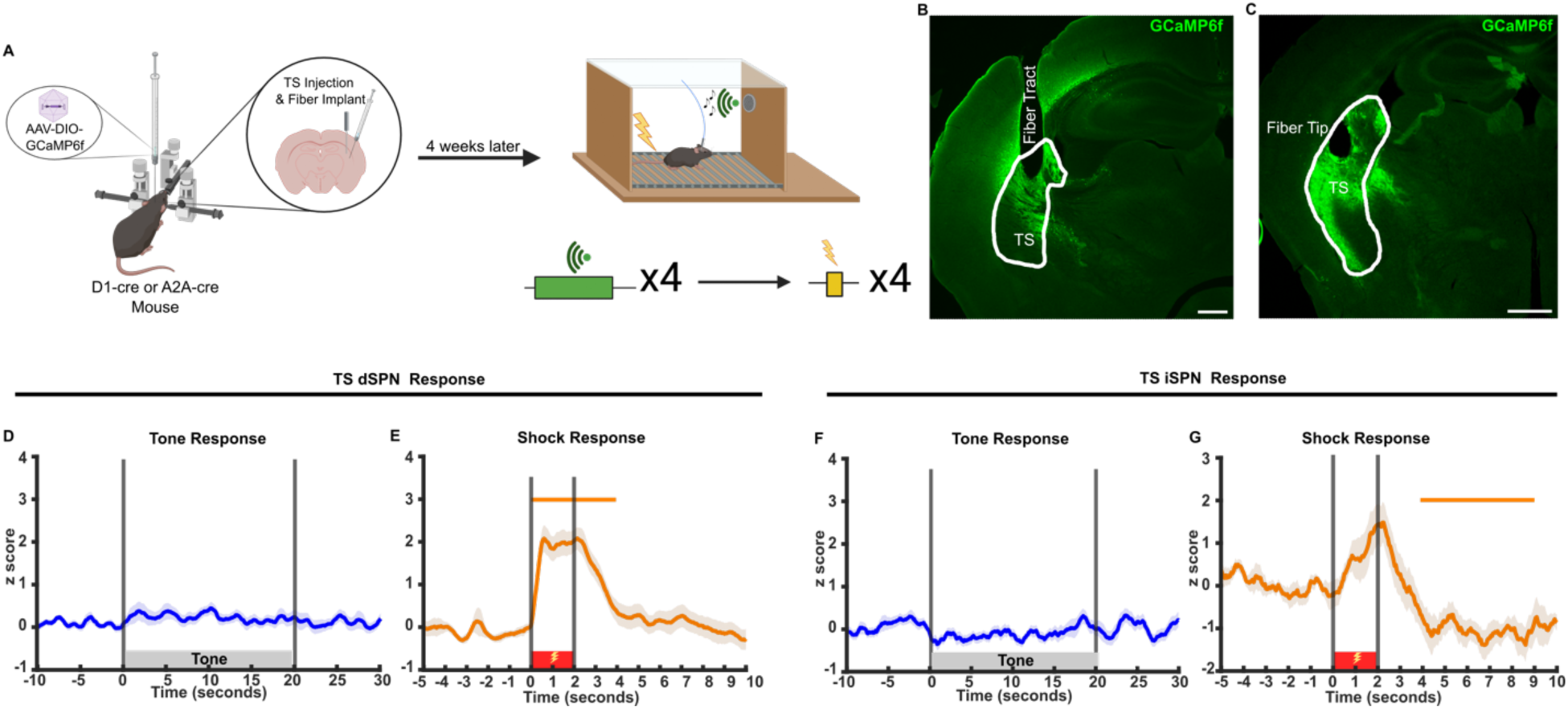
Foot shocks but not neutral sounds evoke responses in TS SPNs. **(A)** Schematic illustration of photometry recording in TS SPNs during sound and foot shock delivery. Created in Biorender. **(B-C)** Representative histology of TS GCamp6f expression and fiber placement in (**B**) D1:cre (dSPN) and (**C**) A2A:cre (iSPN) mice. Scale bar, 400 µm. **(D-E)** TS dSPN calcium responses during **(D)** tone (n=19) and **(E)** shocks (n=9). **(F-G)** TS iSPN calcium responses during **(F)** tone (n=16) and **(G)** shocks (n=8). Bars above transients indicate periods significantly different from a z-score of 0 (p<0.05), and above 0.5 or below -0.5 for each condition.

### Behavioral training

After establishing a lack of TS SPN responses to the tone cue in naive mice, both D1:cre and A2A were assigned to three distinct groups: threat conditioned, safety conditioned and tone alone controls. In *safety-conditioned* mice, the sound cue (CS) was explicitly unpaired from foot shocks during training. *Threat-conditioned* mice received an identical sound cue, but the sounds co-terminated with a foot shock. *Tone-alone control* mice were presented identical sounds with no foot shock (Figure 2A).

**Figure 2.**
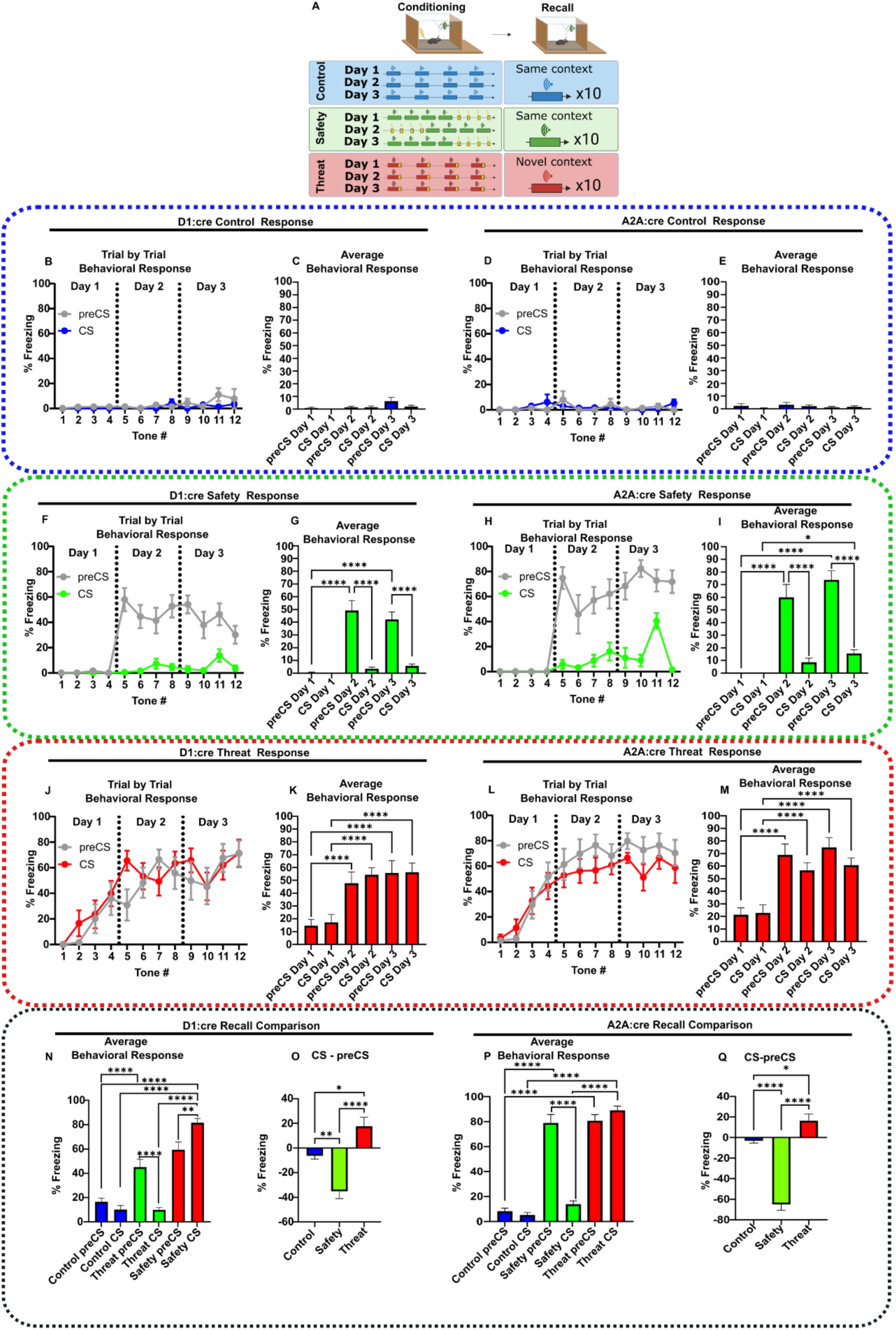
Sound-associative learning in D1:cre and A2A:cre mice. **(A)** Schematic of safety, threat, and control conditioning paradigms. Created in Biorender. **(B-E)** Averaged trial by trial freezing and daily averaged freezing during control conditioning in **(B-C)** D1:cre mice (n=9) and **(D-E)** A2A:cre mice (n=9). **(F-I)** Averaged trial by trial freezing and daily averaged freezing during safety conditioning by **(F-G)** D1:cre (n=10) and **(H-I)** A2A:cre mice (n=8). **(J-M)** Averaged trial by trial freezing and daily averaged freezing during threat conditioning by **(J-K)** D1:cre (n=9) and **(L-M)** A2A:cre mice (n=9). **(N-O)** Average freezing and average difference between preCS and CS freezing (CS-preCS) during recall in control (n=9), threat (n=9) and safety conditioned (n=10) D1:cre mice**. (P-Q)** Average freezing in preCS and CS periods during recall and average difference between preCS and CS freezing (CS-preCS) during recall in control (n=9), threat (n=7) and safety conditioned (n=10) A2A:cre mice. Data represented as mean ± SEM; *p<0.05, **p<0.01, ****p<0.0001. Detailed statistical analysis can be seen in Supplementary Table 2.

As expected, *tone-alone controls* developed no behavioral response (Figure 2B-E).

During *safety-conditioning*, mice did not freeze on training Day 1. However, on Days 2 and 3, mice exhibited extensive freezing in the context (preCS), which was suppressed by CS (Figure 2F-I). The *safety-conditioning* protocol produced a robust learned response to the CS by 24 hours after the initial training, measured as a difference in freezing between preCS and CS. On Day 4, we conducted a recall analysis in which the response to the CS was assessed in the absence of the foot shock. Safety-conditioned mice continued to display context freezing and a robust locomotor response during the CS presentation (Supplementary video 1, Figure 2N-Q, S1A-I).

In contrast, *threat-conditioned mice* developed freezing to the CS on training Day 1, with increased freezing on training Day 2 (Figure 2J-M). There was little difference between preCS and CS responses, indicating that both the context and tones rapidly become aversive. These responses were also retained on the recall test on Day 4 (Supplementary video 2, Figure 2N-Q, S1A-I).

### Safety learning elicited TS SPNs responses to the sound cue

We then analyzed TS SPN activity during training in D1:cre or A2A:cre mice injected in the TS with AAV9-flex-GCaMP6f virus.

In *tone-only controls,* TS SPNs showed no response to the preCS or the CS during training or recall (Figure 3A-C, 4A-C). This suggests that neutral sounds do not activate the TS.

**Figure 3.**
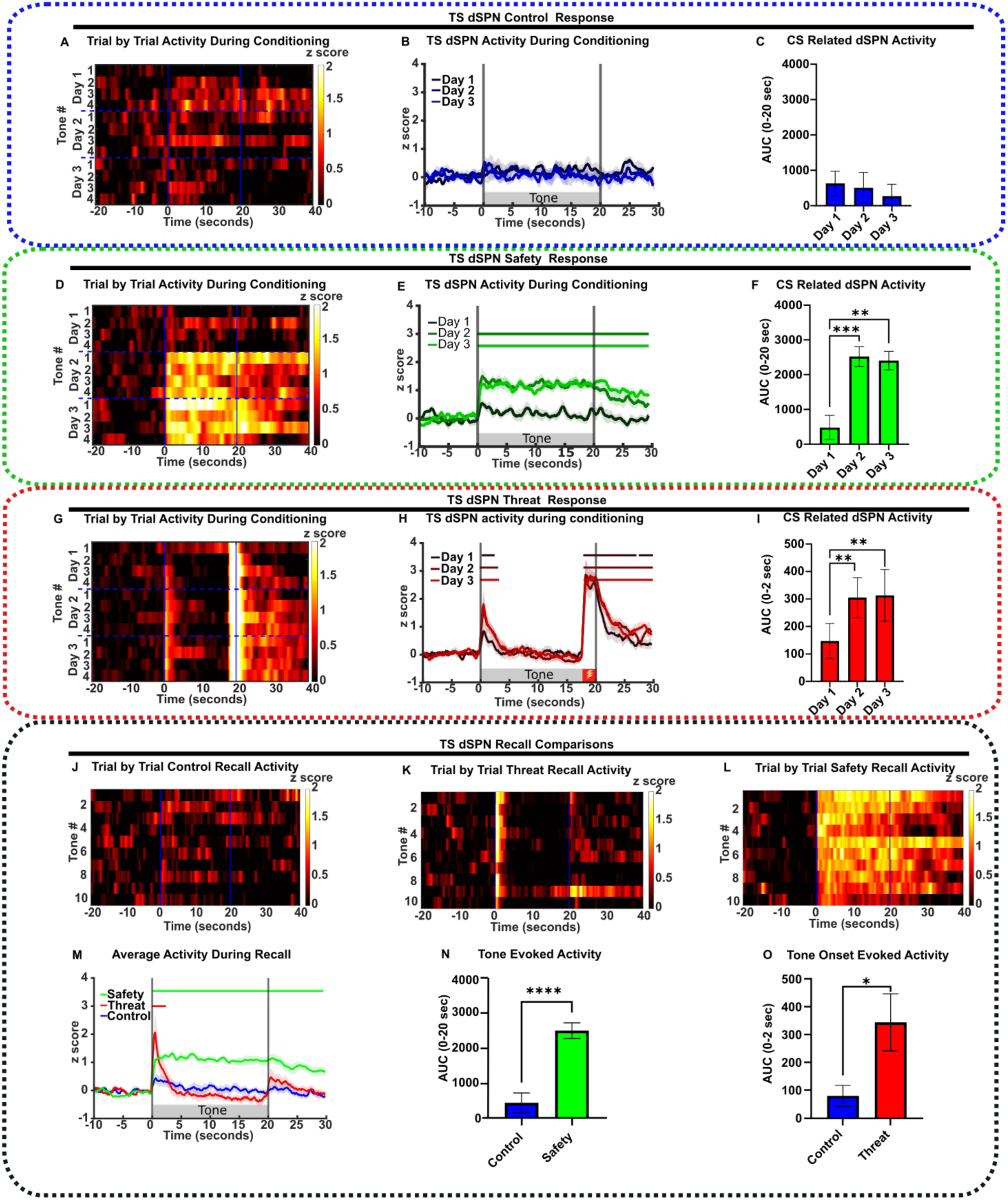
TS dSPN activity correlates with learned safety. (A-I) From left to right, heat map of CS responses across trials, neuronal response during conditioning days and AUC during CS period under **(A-C)** control conditioning (n=9), **(D-F)** safety (n=10) and **(G-I)** threat (n=9). **(J-L)** Heat map of CS responses across **(J)** control recall (n=9)**, (K)** threat recall (n=9) and **(L)** safety recall trials (n=10). **(M)** Average neuronal response during recall for each condition (Safety n=10, Threat n=9, Control n=9). **(N)** Average AUC during the recall trials with CS (0 – 20 seconds) in safety and control conditioned mice (Safety n=10, Control n=9). **(O)** Average AUC at the CS onset (0 – 2 seconds) during recall in threat and control conditioned mice (Threat n=9, Control n=9). All data represented as mean ± SEM; *p<0.05, **p<0.01, ***p<0.001, ****p<0.0001. Bars above transients indicate periods significantly different from z-score of 0 (p<0.05), and above a threshold of 0.5 for each condition. Detailed statistical analysis can be seen in Supplementary Table 2.

**Figure 4.**
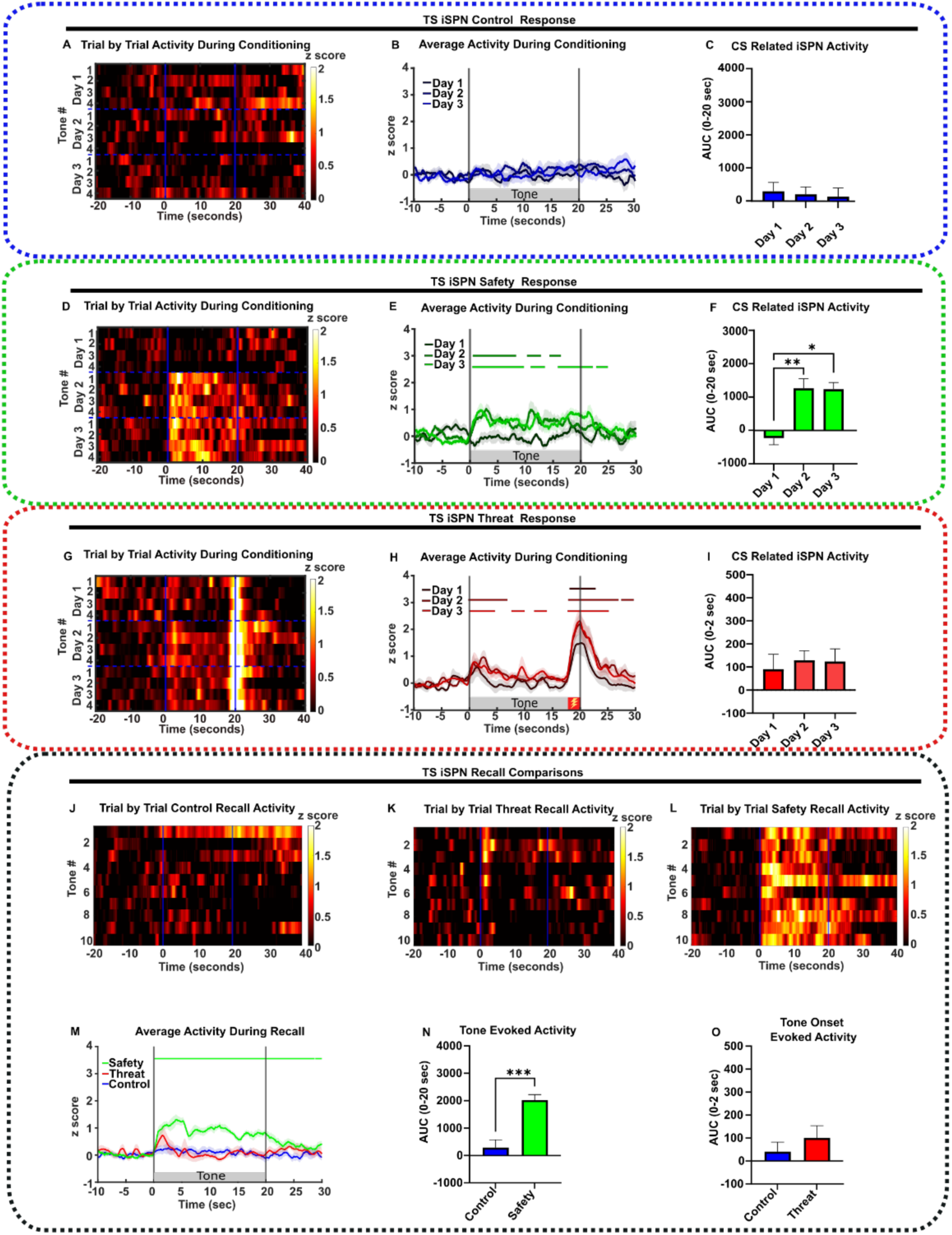
TS iSPN activity correlates with learned safety. (A-I) From left to right, heat map of CS responses across conditioning days, neuronal response, and AUC during the CS period under (**A-C**) control conditioning (n=9), **(D-F)** safety (n=8), **(G-I)** threat (n=9, foot shock indicated by red box). **(J-L)** Heat map of CS responses across **(J)** control recall (n=9), **(K)** threat recall (n=9), and **(K)** safety recall trials (n=10)**. (M)** Average neuronal response during recall (Safety n=8, Threat n=9, Control n=9). **(N)** Average AUC during recall CS presentation (0 – 20 seconds) in safety and control conditioned mice (Safety n=8, Control n=9). **(O)** Mean AUC at the CS onset (0–2 seconds) during recall in threat and control conditioned mice (Threat n=8, Control n=9). All data represented as mean ± SEM; *p<0.05, **p<0.01, ***p<0.001. Bars above transients indicate periods significantly different from z-score of 0 (p<0.05) and above 0.5 for each condition. Detailed statistical analysis is in Supplementary Table 2.

Mice undergoing safety conditioning showed no response to the CS prior to training on Day 1. However, on Day 2, the CS evoked a large increase in activity of both TS SPN populations that lasted throughout the tone duration (Figure 3D-F, 4D-F). These responses were maintained on the recall day (Figure 3J, 3M-N, 4J, 4M-N). The appearance of CS-induced SPN activity on Day 2 was concomitant with the behavioral acquisition of the learned CS responses, and both the neuronal response and behavior were maintained on the recall day when foot shocks were omitted.

These SPN responses differed in *threat-conditioned* mice, as both TS dSPN and iSPN activities exhibited peaks at the onset of the CS (0-2 seconds) that returned to baseline by ∼ 5 seconds (Figure 3M, 4M). The amplitude of the tone onset responses increased on Day 2 of training and was maintained on the recall day (Figure 3G-I, 3M, 3O, 4G-I, 4M, 4O). Interestingly, an additional peak was evoked in dSPNs following shock omission at recall (Figure 3M, S2A-C), which we label an *offset component*. This offset component was absent in iSPNs (Figure 4M, S2D-F).

In addition to tone related responses, foot shock responses were consistently observed in both TS SPN populations in both safety- and threat-conditioned mice (Figure S3).

The results demonstrate that both CS associated behavior and the appearance of TS SPN response to the CS occur by 24 hours after training and are retained during recall.

### The TS dSPN response to CS tone onset remains present under anesthesia

The increased CS-elicited dSPN activity in *safety-conditioning* could be related to a potentiated synaptic response to auditory inputs and/or to activity related to the locomotor response following the tone. To differentiate these components, we measured dSPN and iSPN GCaMP responses to the CS during isoflurane anesthesia, which abolishes motor responses (Figure 5A).

**Figure 5.**
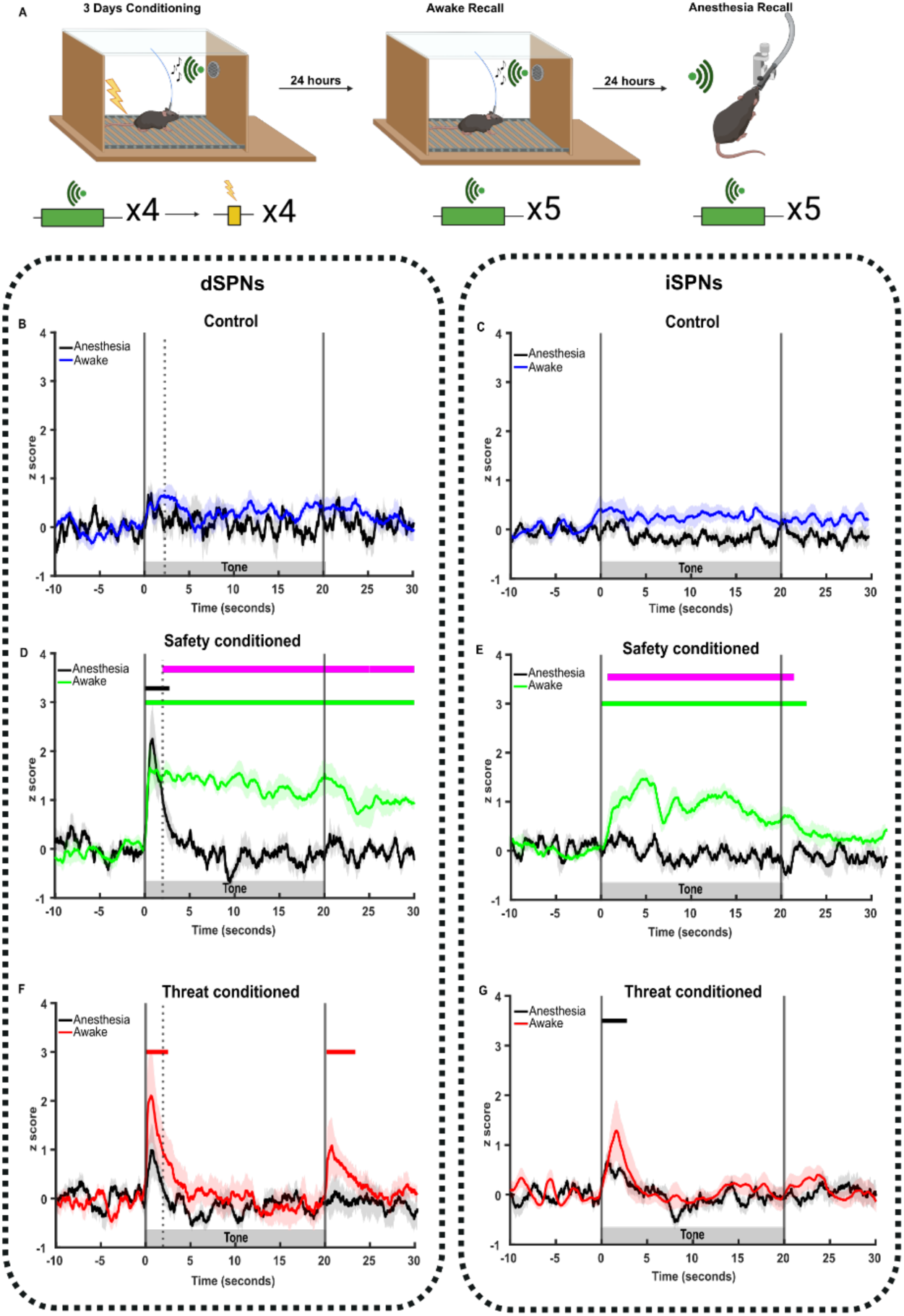
Safety cue induced TS SPN calcium response partially persists under anesthesia. **(A)** Schematic illustrating recall under awake and anesthetized conditions. Created in Biorender. **(B-C)** Control group: average neuronal responses to CS delivered in awake (blue) and anesthetized (grey) conditions in TS dSPN (n=5; B) and TS D2-SPN (n=8; C). **(D-E)** Safety conditioned group: average neuronal responses to CS delivered in awake (blue) and anesthetized (grey) conditions in TS dSPN (n=5; D) and TS iSPN (n=8; E). **(F-G)** Threat conditioned group: average neuronal responses to CS’s delivered in awake (blue) and anesthetized (grey) conditions in TS dSPN (n=5; F) and TS iSPN (n=8; G). Bars above transients show periods significantly different from z-score of 0 for each condition (p<0.05), and magenta bars show periods of significance between conditions.

As expected, in *tone-only control mice*, there was no SPN tone onset component (Figure 5B-C).

In *safety-conditioned mice*, anesthesia abolished the CS-evoked increase in iSPN activity, indicating a requirement for wakefulness. However, the dSPN response to *tone onset* (0-2 sec) in *safety-conditioned mice* was retained during anesthesia, while the *sustained component* (2-20 sec) was abolished (Figure 5D-E).

In *threat-conditioned mice*, anesthesia decreased the CS onset transient in both dSPNs and iSPNs. Anesthesia also abolished the offset component that is particular to the dSPNs when foot shock is omitted, indicating a requirement for consciousness and/or movement for that response (Figure 5F-G).

As a control to confirm if the dSPN response to the safety cue was related to neuronal activity, we administered the neuronal activity blocker tetrodotoxin (TTX) via a unilaterally implanted optofluid cannula in the TS (Figure 6A). As expected, TTX blocked all GCaMP responses to CS and foot shocks in awake mice (Figure 6C-E), indicating that the responses above are not movement artifacts. Moreover, the behavioral responses to CS and context were reduced by TTX infusion (Figure 6F-G), consistent with a role of the TS in context-evoked threat and tone-evoked safety responses.

**Figure 6.**
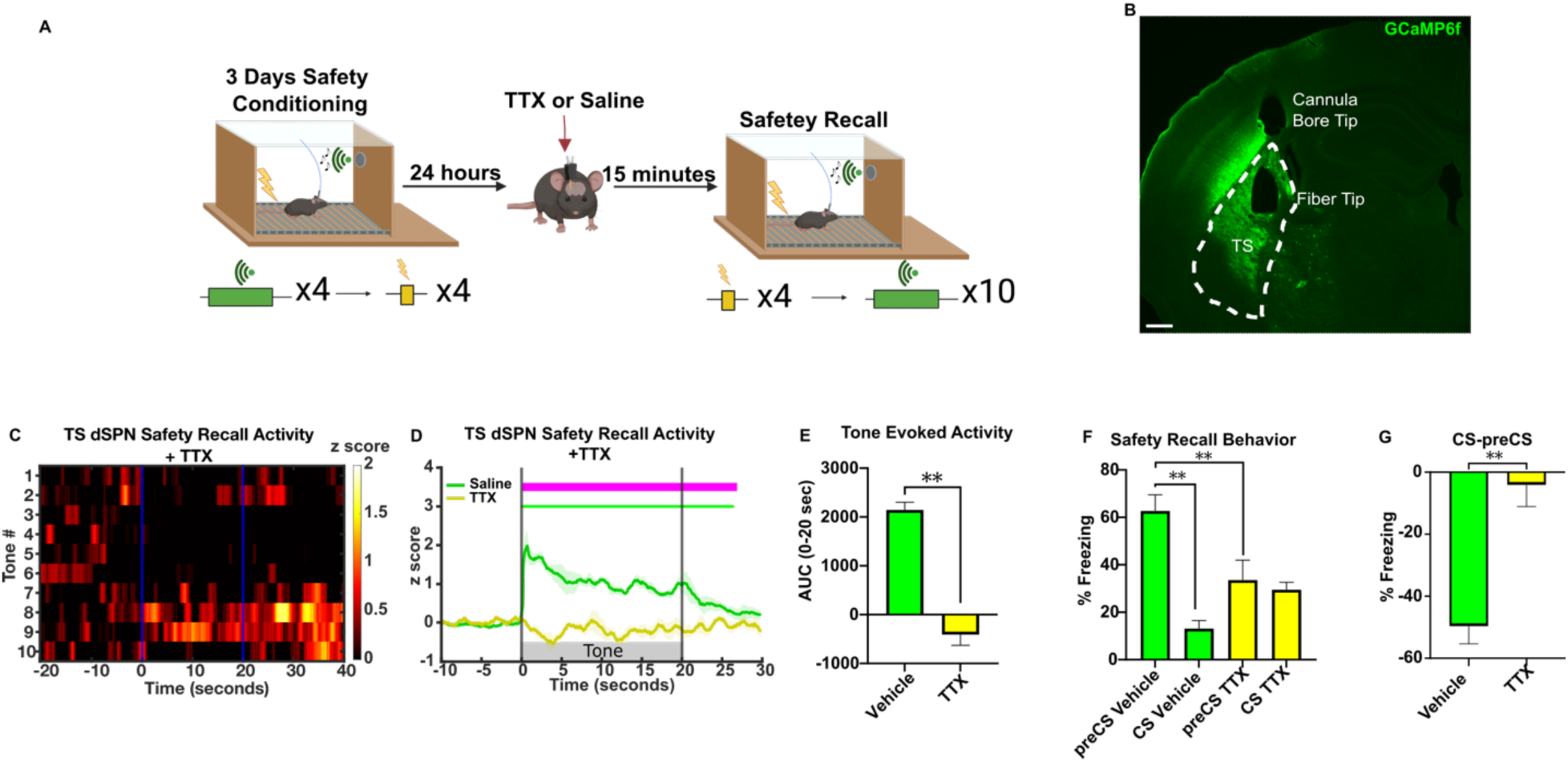
TS inactivation attenuates learned safety responses. **(A)** Schematic illustration of safety conditioning followed by optofluidic drug infusions prior to shock and safety recall. Created in Biorender. **(B)** Representative histology of TS GCamp6f expression and fiber placement in D1:cre mice. Scale bar 400 µm. **(C)** Heatmap of average safety CS response across recall trials under TTX activity blockade (n=5). **(D)** Safety cue induced TS dSPN calcium response following vehicle or TTX administrations. Bars above transients show periods significantly different from z-score of 0 (p<0.05) and above 0.5 for each condition, and magenta bar indicates periods of significance between conditions (n=5). **(E)** Average AUC during recall following vehicle or TTX administration (n=5). **(F)** Average percent time freezing in preCS and CS periods during safety recall following TTX inactivation of TS (n=5). **(G)** Percent time freezing difference (CS-preCS) during recall following TTX administration in TS (n=5). Detailed statistical analysis is in Supplementary Table 2.

We then performed a correlation analysis of CS-elicited dSPN activity relative to locomotion (Figure S4). We found that the dSPN *tone onset-locked component* activity was not correlated with movement during recall (Figure S4E and S4G), while the *sustained component* was movement-related (Figure S4F and S4H), consistent with the blockade of the sustained component by anesthesia.

Together, these results indicate that safety conditioning produces a potentiated anesthesia-independent response to auditory inputs at CS *tone onset* in dSPNs.

### Sound-safety associative learning doesn’t alter dSPN intrinsic excitability

We then analyzed the mechanisms by which the CS produced the learning-dependent tone onset response by TS dSPN neurons. To identify these neurons for electrophysiological recordings, we injected D1:cre mice in the TS with a cre-dependent tdTomato virus. After four weeks of viral expression, mice were *safety conditioned* or untreated. On Day 4, corresponding to the recall day in the in vivo experiments, the mice were sacrificed and tdTomato+ dSPNs in the TS were analyzed by whole cell patch clamp (Figure 7A).

**Figure 7.**
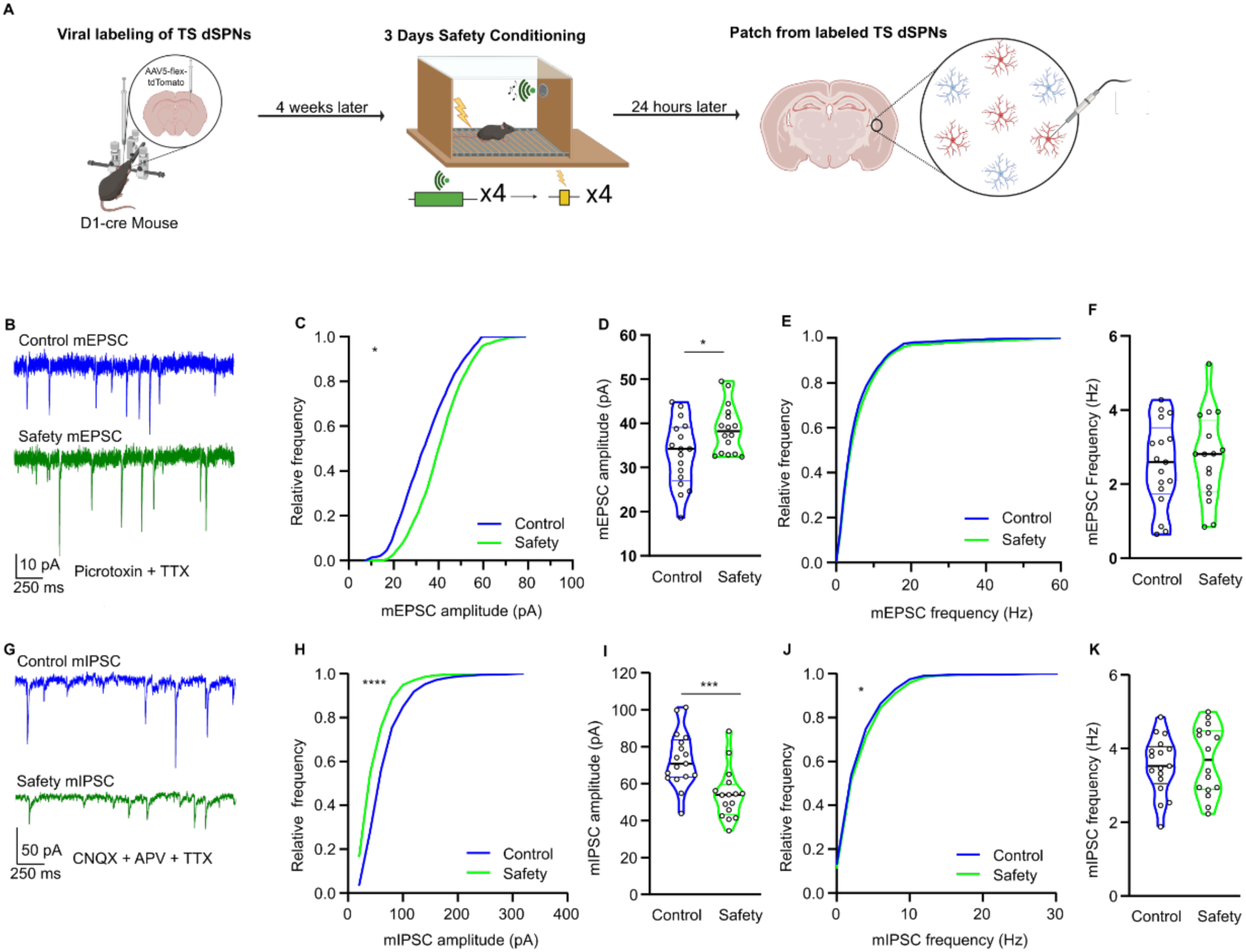
TS dSPNs of safety trained mice are intrinsically less excitable. **(A)** Schematic illustrating viral labeling, safety training, and patch clamping strategy. Created in Biorender. **(B)** Representative traces of miniature EPSCs (mEPSCs) from control and safety trained groups in the presence of 25 µM picrotoxin and 1 µM TTX. **(C)** Cumulative histogram for mEPSC amplitude (Kolmogorov-Smirnov test, p=0.0142, Control: N=7, n=15, Safety: N=6, n=16). **(D)** Average mEPSC amplitudes. (Mann-Whitney, p=0.0445, Control: N=7, n=17, Safety: N=6, n=16). **(E)** Cumulative histogram for mEPSC instantaneous frequency (Kolmogorov-Smirnov test, p= 0.0805, Control: N=7, n=15, Safety: N=6, n=16). **(F)** Average frequency of mEPSC (Mann-Whitney test, p=0.7356, Control: N=7, n=17, Safety: N=6, n=16). **(G)** Representative traces of miniature IPSCs (mIPSCs) from control and safety trained groups in the presence of 25 µM APV, 10 µM CNQX, and 1 µM TTX. **(H)** Cumulative histogram for mIPSC amplitude (Kolmogorov-Smirnov test, p<0.0001, Control: N=6, n=17, Safety: N=5, n=16). **(I)** Average mIPSC amplitudes (Mann-Whitney t test, p=0.005, Control: N=7, n=17, Safety: N=6, n=16). **(J)** Cumulative histogram for mIPSC instantaneous frequency (Kolmogorov-Smirnov test, p=0.0487, Control: N=7, n=17, Safety: N=6, n=16) **(K)**. Average frequency of mIPSC (Mann-Whitney t test, p=0.7090, Control: N=7, n=17, Safety: N=6, n=16). The violin plots are displayed as median ± quartiles. All other data was represented as mean ± SEM; *p<0.05, ***p<0.001. Detailed statistical analysis can be seen in Supplementary Table 2.

We first examined whether the enhanced TS dSPN response to the onset of auditory cues is related to enhanced intrinsic excitability. Measures of intrinsic membrane properties, including resting membrane potential, input resistance, and membrane capacitance, were unaltered by safety learning (Figure S5A-D). Consistently, sound safety association did not affect the excitability of TS dSPN, as shown by identical responses to depolarizing step currents (-10 to +390 pA in 20 pA increments at 500 ms duration) (Figure S5E) and current ramps (0-800 pA; 200 pA/s) (Figure S5G) in conditioned animals and controls. The minimum current threshold required to elicit the first action potential (rheobase) was not altered by sound-safety association (Figure S5F). We did not observe significant changes in voltage-gated ion currents known to control the excitability of SPNs, including voltage-gated Ca^2+^ (*30*) (Figure S5H-I), inward rectifying potassium (Kir) (*31*) (Figure S5J-K), and A-type potassium (K_a_) (*32*) currents (Figure S5L-M). Together these results indicate that sound-cued safety learning did not alter the intrinsic excitability of TS dSPNs.

### Safety learning enhances postsynaptic TS dSPN responses to excitation

An alternative hypothesis for the enhanced TS dSPNs response to CS, is an increased synaptic response to glutamatergic inputs from the cortex and thalamus(*33*). To analyze whether sound-safety association altered synaptic response to excitatory input, we measured spontaneous miniature excitatory postsynaptic currents (mEPSCs) in the presence of TTX in TS slices from safety conditioned and control mice. The amplitude of mEPSCs in safety conditioned mice was significantly higher than controls, while the mEPSC frequency was unchanged (Figure 7B-F), indicating that safety conditioning enhanced the postsynaptic but not presynaptic responses to excitatory inputs. We then examined inhibitory miniature inhibitory postsynaptic currents (mIPSCs) and found that safety conditioning reduced the amplitude of mIPSCs but not their frequency (Figure 7G-K). Together, these results indicate that safety training enhanced the postsynaptic response by dSPNs to glutamatergic synapses and decreased their response to GABAergic synapses.

To examine the basis for enhanced dSPN responses to excitatory synaptic inputs, we electrically stimulated the corpus callosum, which provides cortical input throughout the striatum (Figure S6A) in the presence of the GABA_A_R antagonist picrotoxin. We found that sound safety association enhanced the corticostriatal synaptic responses (Figure S6B).

We then examined responses to inhibitory synaptic inputs, which originate from local inhibitory interneurons and long range inhibitory cortical projections (*34–36*) by electrically stimulating the TS in the presence of glutamate antagonists (CNQX and APV). The response to these inhibitory inputs was dampened by sound safety association (Figure S6F). Paired pulse stimulation revealed no difference between control and safety conditioned mice (Figure S6G). These results are consistent with the mEPSC and mIPSC data and confirm that sound-safety association elicits changes in synaptic plasticity via postsynaptic mechanisms.

To examine dSPN responses to auditory thalamic synaptic inputs, which send axonal inputs to both cortex and striatum (*37–39*), we injected a cre-independent Chronos virus into the auditory thalamus and a cre-dependent tdTomato virus into the TS of D1:cre mice (Figure 8A). Auditory thalamic axons within the TS were optically stimulated (470 nm, 5 ms) to induce optical EPSCs (oEPSCs) (Figure 8B-D). Safety conditioning enhanced the dSPN responses to oEPSCs (Figure 8B). Presynaptic plasticity was unchanged as indicated by similar paired-pulse ratios between safety-trained and control mice (Figure 8C). These findings are also consistent with the mEPSC results and further confirm that safety conditioning strengthens postsynaptic responses without inducing presynaptic plasticity.

**Figure 8.**
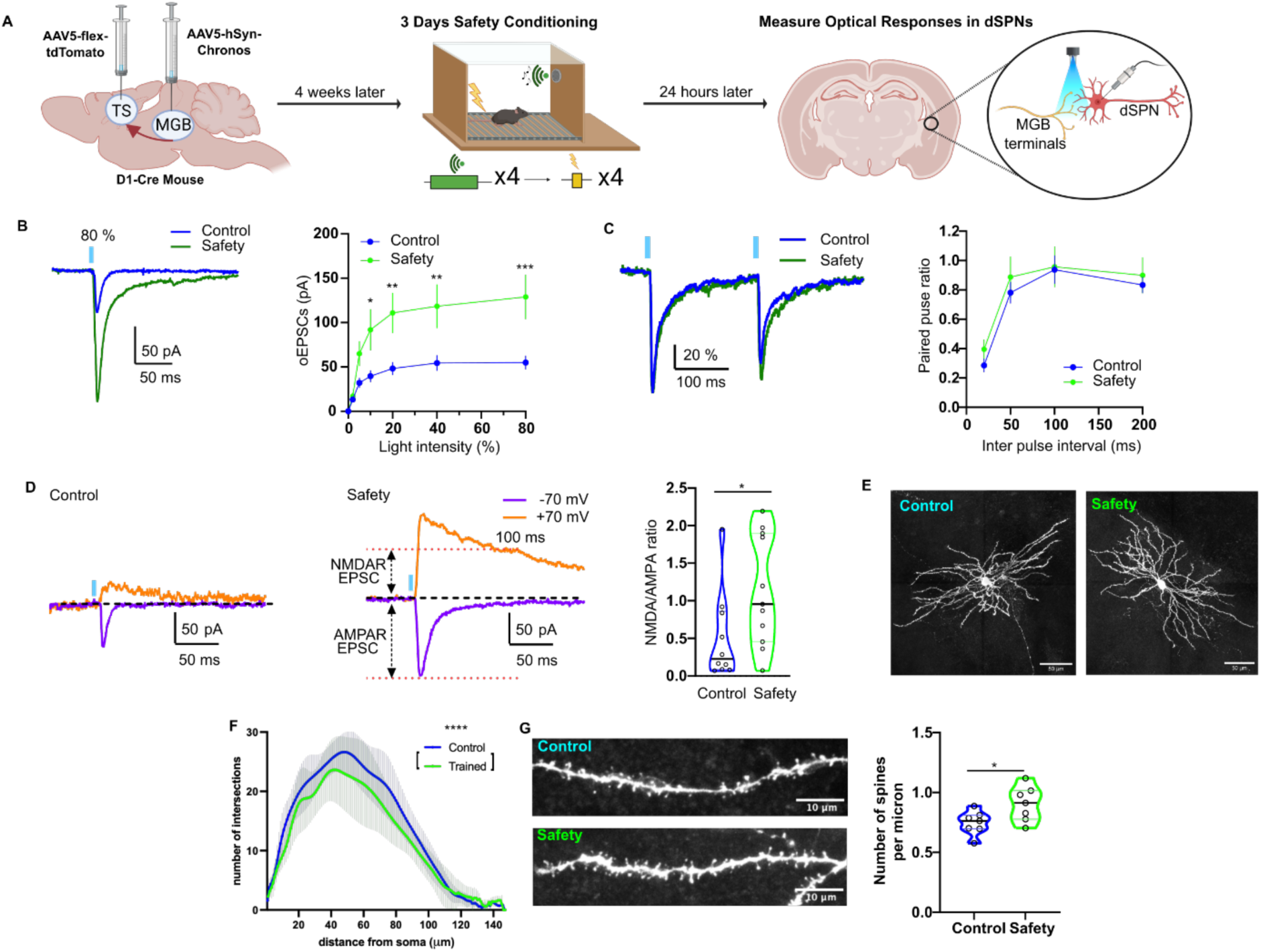
TS dSPNs of safety trained mice are more responsive to auditory thalamic inputs. (A) Schematic illustrating viral, safety training, and patch clamp strategy. Created in Biorender. (**B**) Left: Representative traces of optical EPSCs (oEPSCs) evoked by auditory thalamic projections (60 % light intensity) from control and safety trained groups. Right: Thalamo-striatal synaptic transmission in response to increasing light intensities (5 ms duration; 0, 2, 5, 10, 20, 40, and 80 %) in safety conditioned group (Two-way ANOVA followed by Bonferroni’s test, light intensity factor (F(6, 224) = 16.91 p<0.0001), group factor (F(1, 224) = 39.32 p<0.0001), interaction factor (F(6, 224) = 2.928, p=0.009), Control: N=7, n=20, Safety: N=5, n=14)**. (C)** Left: Representative traces of paired oEPSCs evoked by two identical stimuli at 200 ms interval. The traces were normalized to maximal amplitude of the first oEPSCs respectively. Right: Paired-pulse ratios over a range of inter-pulse intervals (Control: n=14, Safety: n=13). **(D)** Representative AMPAR and NMDAR oEPSCs to optical stimulation in TS dSPNs from control (left) and safety conditioned (middle) mice. Right: Quantification of AMPAR/NMDAR ratio at +70 mV (Two-way ANOVA followed by Bonferroni’s test, Inter pulse interval factor (F(3, 170) = 16.34 p<0.0001), group factor (F(1, 170) = 1.215 p=0.2718), interaction factor (F(3, 170) = 0.09355, p=0.9635), Control: N=9, n=24, Safety: N=8, n=21). **(E)** Representative biocytin-filled TS dSPNs from control (left) and safety conditioned (right) mice. **(F)** Number of dendritic branching over various distances from the soma (Control: N=6, n=8, Safety: N=4, n=7). **(G)** Left: Representative dendrites for control and safety conditioned mice. Right: Number of spines per micron averaged per cell (Control: n=7, Safety: n=7). The violin plots are displayed as median ± quartiles. All other data was represented as mean ± SEM; *p<0.05, **p<0.01, ***p<0.001, ****p<0.0001. Detailed statistical analysis can be seen in Supplementary Table 2.

We further characterized the enhanced dSPN response by measuring the NMDAR and AMPAR components of the responses to auditory thalamic input. We found that both components were enhanced by safety conditioning (Figure S6C), with an increased NMDAR to AMPAR ratio (Figure 8D).

We then examined if the enhanced thalamostriatal response might be related to decreased long-term depression (LTD) (*40*). The induction and maintenance of LTD at thalamostriatal dSPN synapses by low frequency (10 Hz) thalamic stimulation paired with postsynaptic depolarization was identical in control and safety conditioned mice (Figure S5D-E).

In contrast to other regions in the striatum, in the TS, auditory cortical and auditory thalamic inputs both converge onto SPN dendritic spines (*41*). To determine if the enhanced responses by TS dSPN glutamate receptors was correlated with morphological alterations, we examined biocytin labelled TS dSPNs. Scholl analysis revealed that while there were fewer dendritic branching intersections in TS dSPNS of safety conditioned mice that there was a higher density of dendritic spines than in controls (Figure 8E-G, S6H). No significant difference was observed in dendritic length, number of primary dendrites, soma surface area or number of branch points (Figure S6I-L).

## DISCUSSION

The learned association of a sound with safety has been a topic of interest since foundational studies by Ivan Pavlov (*3, 11–14, 42*). However, the synaptic changes responsible for these processes have been unknown. We report that safety learning, a form of auditory associative learning, produces powerfully enhanced TS SPN responses to a CS that occurs by the second day of training, and that this is concurrent with the learned behavioral response. The learning-induced synaptic modifications consist of a sound cue onset component in dSPNs that remains during anesthesia and is uncorrelated with movement, followed by a sustained components that is blocked by anesthesia and is associated with locomotion.

We find that this enhanced TS dSPN activity during the onset of CS tone is associated with postsynaptic changes that increase responses to excitatory synapses and decrease responses to inhibitory synapses. It is striking that, as the tone onset-locked component of learned TS dSPN sound response persisted even under anesthesia, the postsynaptic changes induced by auditory associative learning learned remain during unconscious states, despite the absence of locomotion. Previous reports indicate that cortical inputs are required for discrimination between distinct cues (*43*), while safety and threat learning depend on the auditory thalamus and not the auditory cortex (*44, 45*). As isoflurane anesthesia decreases overall cortical activity more than thalamic activity (*39, 46–51*), it may be that the response to thalamic inputs provides the CS anesthesia-resistant tone onset-locked component.

We observed that auditory learning produced a strong potentiation of both NMDA and AMPA components on auditory thalamus-TS dSPN synapses. In addition to calcium-permeable NMDA receptors, SPNs express Ca^2+^-permeable AMPA receptors that contain the GluA1 subunit (*52–54*). Therefore, both AMPA and NMDA activation at auditory thalamo-TS dSPN synapses would generate Ca^2+^ influx during safety learning. This may facilitate Ca^2+^ dependent processes including state transitions (*55*), local dendritic spike modulation (*56*), or the induction of new synapses. Distinguishing these mechanisms are goals for further investigation.

How would enhanced postsynaptic responses by TS dSPN responses underlie learned behaviors? Recently, a direct projection from TS SPNs to the lateral amygdala, a region critical to sensory outcome associations, was shown to regulate threat associative learning (*16*), although its role in sound-safety association was not reported to our knowledge. TS dSPNs are GABAergic neurons that project to the lateral SNr, which in turn sends GABAergic projections to multiple nuclei that control motor output including the periaqueductal grey (PAG), superior colliculus (SC), inferior colliculus (IC), and ventral anterior thalamic nucleus (VAmc) (*57–60*). Freezing is controlled by the PAG and modulated via projections from the SC and IC (*61, 62*). The SC and IC are also implicated in auditory perception, sensory cue elicited movements (*63, 64*), and defensive behaviors (*58, 62, 65*). The VAmc regulates motor activity via excitatory projections to the pre-supplementary motor cortex (preSMA) (*66–69*). Therefore, activation of TS dSPNs by learned safety cues could promote locomotor activity by disinhibiting the VAmc and exciting the preSMA. In contrast, TS iSPNs are GABAergic neurons that project indirectly to the lateral SNr via the GPe and subthalamic nucleus (STN) and disinhibit the SNr, and so act to inhibit the VAmc and the preSMA (*67, 70, 71*).

During locomotion, both dSPNs and iSPNs are active in the anterior dorsal striatum (also known as the motor striatum), where dSPNs are modeled to promote desired movements while iSPNs inhibit unwanted movements (*15, 72*). The SPNs in the TS may serve an analogous function in modulating freezing or locomotor activity in a sensory-dependent manner. In support of this hypothesis, TS D1 SPNs promote while TS D2 SPNs suppress threat avoidance (*73*). Alternatively, TS dSPNs may enhance and iSPNs may inhibit sensory cortices, analogous to the role of anterior dorsal striatal projections to motor cortices (*74*). A prefrontal cortex (PFC)-TS-GPe-thalamic reticular nucleus (TRN)-auditory thalamus pathway is reported to be critical for suppressing attention to irrelevant sensory stimuli (*75*), suggesting that TS dSPNs promote and TS iSPNs inhibit sensory attention (*76–80*). Such a role is consistent with reports that optogenetic stimulation of TS dSPNs or auditory inputs to the TS biases movements contralateral to the side of stimulation in an auditory guided decision task (*77–81*).

Together, these results indicate that auditory learning-driven TS synaptic plasticity consists of increased excitatory and decreased inhibitory postsynaptic response in dSPNs to an associated sound. The identification of this plasticity may suggest treatments for disorders with impaired safety processing, including PTSD (*6, 7, 82*) and Parkinson’s disease (*83*), and suggest implications for synaptic mechanisms that could underlie other forms of sound associative learning, including language acquisition, in which TS activity is implicated (*84*).

## Methods

### EXPERIMENTAL MODELS AND SUBJECT DETAILS

All experimental procedures were approved by the Columbia University Institutional Animal Care and Use Committee (IACUC). All mice in this study were on a C57BL/6J background, typically group housed with littermates of the same sex, and given access to food and water ad libitum. Mice were kept on a reverse light cycle (lights on from 10:00 PM to 10:00 AM) and testing was performed during the dark period under red light. Adult male and female mice between 2 and 12 months of age (6 months average, see Figure S1L) were used for photometry experiments. The following mouse lines were used in this study: D1:Cre (strain 030989-UCD) sourced from Mutant Mouse Resource & Research Centers and A2A:Cre (strain 010687) sourced from The Jackson Laboratories. These mouse strains have been validated for selective expression in dSPNs and iSPNs respectively (*26, 29*). Heterozygous D1:Cre and A2A:Cre mice were used for all experiments and were maintained by backcrossing to C57/B6J wild-types. Experimental groups contained similar numbers of male and female mice. No sex differences were observed in learned safety CS responses, although subtle sex differences in contextual threat responses were observed in D1:cre mice (Supplementary Table 1).

## METHOD DETAILS

All schematics were created in Biorender.

### Surgical procedures

Injection and implant surgeries were performed under 2% isoflurane anesthesia using a stereotaxic instrument (Kopf instruments). A small incision was made in the scalp and burr holes were drilled in the skill at the appropriate stereotaxic coordinates (AP, ML, and DV relative to bregma): -1.06 AP, -3 ML, -3.25 DV for TS; -3.2 AP, -2 ML, -3.8 DV for MGB. Viruses were infused using a glass micropipette, backfilled with mineral oil, connected to microinjector (Nanoject II Drummond Scientific Company) at a rate of 23 nl/ min. The injection pipette was slowly withdrawn 5 minutes after the end of infusion. For fiber photometry experiments, optical fibers (Doric Lenses, 0.22 NA, 4 mm length, 300 µm diameter flat tip) were implanted 0.1mm above the target injection site. For optofluidic photometry experiments, optofluidic cannula’s (Doric Lenses, 0.37 NA, 5mm fiber length, 300 µm diameter flat tip, 4.9 mm cannula bore length) were implanted 0.1mm above target injection site.

### Viral volumes, sources and titers

For fiber photometry experiments targeting the TS, 230 nl of AAV9-Syn-Flex-GCaMP6f-WPRE-SV40 (addgene,1*10^13^ vg/ml) was unilaterally injected at a rate of 23 nl per minute. An optical fiber (Doric Lenses; MFC 300/370-0.22_4 mm_MF2.5_FLT) or optofluid cannula (Doric Lenses; OmFC_MF1.25_300/370-0.22_3.2_FLT_3) was implanted 0.1 mm above the target site. For electrophysiology experiments, AAV5-Syn-Chronos-GFP (addgene,1*10^13^ vg/ml) was injected in the MGB and AAV9-Flex-tdTomato (addgene,1*10^13^ vg/ml) was injected in the TS. Mice were given 4 weeks for surgery recovery and viral expression before conducting experiments.

### General histology procedures and imaging

Mice were deeply anesthetized and transcardially perfused with 0.9% NaCl followed by 4% paraformaldehyde (PFA) in 0.1 M phosphate buffer (PB). Brains were removed, post-fixed overnight in 4% PFA in 0.1M PB, sectioned coronally at room temperature on a VT1200 vibratome (Leica Biosystems) and stored in cryoprotectant (0.1 M PB, 30% glycerol, 30% ethylene glycol) at −20°C. For immunofluorescence analysis, 40 µm sections were washed in TBS for 10 minutes three times and then blocked and permeabilized for 1 h at room temperature with 10% normal donkey serum (Jackson ImmunoResearch) and 0.1% Triton-X in TBS. Sections were then incubated overnight at 4°C with primary antibody (Chicken anti-Green Fluorescent Protein; 1:500; ab13970, Abcam) for GCaMP staining in 2% normal donkey serum, 0.1% Triton-X in TBS. Sections were then washed for 10 minutes in TBS three times and incubated in secondary antibody (Goat anti-Chicken Alexa Fluor 488; 1:500; A-11039, Invitrogen) for 1 hour. Sections were then washed for 10 minutes in TBS three times and mounted on microscope slides. Images were obtained using an epifluorescence microscope (Olympus IX81 Microscope, Boston Industries, Inc. Walpole, MA) equipped with 10X objective lens.

### Slice preparation

Whole-cell recordings were performed on 4–6-month-old D1:cre mice. These mice underwent stereotaxic injection of cre independent Chronos virus in the MGB and cre dependent tdTomato virus in the TS to allow for identification of dSPNs as well as selective stimulation of auditory thalamic terminals. After 4-5 weeks to allow for viral expression, mice were safety conditioned for 3 days and were euthanized by cervical dislocation on day 4. These mice were decapitated and coronal 250 µm-thick slices were prepared on a vibratome (VT1200; Leica, Sloms, Germany) in oxygenated ice-cold cutting-artificial cerebrospinal fluid (ACSF) containing 194 mM sucrose, 30 mM NaCl, 4.5 mM KCl, 26 mM NaHCO_3_, 6 mM MgCl_2_·6H_2_O, 1.2 mM NaH_2_PO_4_, and 10 mM D-glucose (pH 7.4, 290 ± 5 mOsm). Slices containing the TS, indicated by the presence of the basolateral amygdala, were collected and transferred to oxygenated normal ACSF containing 125.2 mM NaCl, 2.5 mM KCl, 26 mM NaHCO_3_, 1.3 mM MgCl_2_·6 mM H_2_O, 2.4 mM CaCl_2_, 0.3 mM NaH_2_PO_4_, 0.3 mM KH_2_PO_4_, and 10 mM D-glucose (pH 7.4, 290 ± 5 mOsm) at 34 °C and allowed to recover for at least 40 min before electrophysiological recordings.

### Whole-cell patch clamp recording

The recording chamber was continuously perfused with normal ACSF (1.5-2 ml/min) at 34°C. Electrophysiological recordings were performed on an upright Olympus BX50WI differential interference contrast microscope (Olympus, Tokyo, Japan) equipped with a 40 x water immersion objective and an infrared video camera. Visualization of TS dSPNs expressing tdTomato was achieved with 555 nm light illumination. Patch pipettes (5-6 MΩ) were pulled using P-97 puller (Sutter instruments, Novato, CA) and filled with solution indicated below. Whole-cell patch clamp recordings were performed with a MultiClamp 700B amplifier (Molecular Devices, Forster City, CA) and digitized at 10 kHz with InstruTECH ITC-18 (HEKA, Holliston, MA).

Intrinsic membrane properties were measured in current clamp mode using a pipette solution containing 115 mM K-gluconate, 10 mM HEPES, 2 mM MgCl_2_, 20 mM KCl, 2 mM MgATP, 1 mM Na_2_-ATP, and 0.3 mM GTP (pH=7.3; 280 ± 5 mOsm). To induce current-voltage responses, 500 ms duration somatic step currents were injected through the patch pipette. Rheobase was obtained by ramp current injection from 0-800 pA at 200 pA/s. In voltage clamp mode, glutamatergic synaptic transmission was measured at -70 mV holding potential in the presence of 25 µM picrotoxin with a pipette solution containing 120 mM CsMeSO_3_, 5 mM NaCl, 10 mM HEPES, 1.1 mM EGTA, 2 mM Mg^2+^-ATP, 0.3 mM Na-GTP, 2 mM Na-ATP, 10 mM TEA-Cl (pH=7.3, 280 ± 5 mOsm). Miniature excitatory postsynaptic currents (mEPSCs) were detected in the presence of (1 µM TTX). To induce evoked corticostriatal synaptic transmission, concentric tungsten electrode (WPI, Sarasota, FL) was placed on the corpus callosum and electrical stimuli were delivered by Master-9 (A.M.P.I., Jerusalem, Israel) with stimulation intensity adjusted by Iso-Flex (A.M.P.I.). Optical stimulation (470 nm) of auditory thalamic terminals was achieved by LED illumination (Colibri 2, Zeiss, Germany) through 40x objective at 10 - 80 % of light intensity controlled by Master-9. To measure NMDA components, cells were held at +70 mV and currents were measured 100 ms after the time of optical stimulation. 25 µM of the NMDA antagonist APV was bath applied to pharmacologically isolate AMPA currents. Long-term depression of auditory thalamus to TS dSPN synapse was induced by 10 Hz presynaptic stimulation (3000 pulses for 5 min) paired with +70 mV postsynaptic depolarization. Voltage gated ion channel currents were examined in the presence of TTX (1 µM) and synaptic blockers. Inward-rectifying (Kir) and A type potassium currents (K_a_) were measured using K-gluconate pipette solution. Kir currents were induced by 1 s step voltages (V_h_= -70 mV -150 - -50 mV +10 mV increments), and K_a_ currents were measured by 0.5 s step voltages (V_h_= -100 mV, 0 - +120 mV, +10 mV increments). CsMeSO_3_ pipette solution was used to measure voltage-gated Ca^2+^ currents with 100 ms step voltages (V_h_= -110 mV, -70 - +40 mV, +10 mV increments). Data was sampled at 20 kHz and filtered at 4 kHz. The ion currents were analyzed after leak subtraction using P/4 protocol. All chemicals for ACSF, pipette solution, and reagents were purchased from Sigma-Aldrich and Tocris. GABAergic miniature inhibitory postsynaptic currents (mIPSCs) were recorded in the presence of 25 µM APV, 10 µM CNQX, and 1 µM TTX. Cells were held at -70 mV with pipettes containing 140 mM CsCl_2_, 2 mM MgCl_2_, 10 mM HEPES, 2 mM EGTA, 2 mM MgATP, 1 mM Na_2_-ATP, 0.3 mM GTP and 5 mM QX314 (pH = 7.3, 280 ± 5 mOsm). To induce evoked IPSCs, the stimulus electrode was placed ∼100 µm away from the recorded neurons.

Data was acquired using WINWCP software (developed by John Dempster, University of Strathclyde, UK) and analyzed using Clampfit (Molecular Devices), Igor Pro (Wavemetrics, Lake Oswego, OR), and MATLAB (MathWorks, Natick, MA) and graphed with Prism software (GraphPad Software Inc, CA). The violin plots are displayed as median ± quartiles and line graphs are expressed as mean ± standard error of the mean (SEM). Data in the text are expressed as mean ± SEM. Kolmogorov-Smirnov test was used for group comparison of distribution of miniature events (mEPSC and mIPSC). The statistical tests used are listed in the figure legends and text. Unless indicated, two-way ANOVA and non-parametric unpaired t-tests were used to compare groups for all figures

### Biocytin diffusion and morphological analysis

Direct pathway SPNs were identified by somatic tdTomato expression and whole-cell patch clamp recording was performed. One cell per slice was filled with 0.1-0.2 % of biocytin for at least 15 minutes. Slices were then fixed in 4 % paraformaldehyde for 24 hours and transferred to phosphate buffered saline (PBS). After three washes in PBS, slices were transferred to Tween - tris buffered saline (TTBS) (0.6 %) with streptavidin conjugated with Alexa Fluor 647 or DyLight 649 (1:200) and incubated for 72 hours. Mounted slices were dried overnight and then a cover slip with an anti-fading mounting medium (Fluoromount G) was applied. Images were acquired using the Leica SP8 confocal microscope with a 63x/1.4 NA Zeiss lens. Whole-cell images were obtained at 0.15 µm pixel resolution with 0.5 µm z-steps. For planar Sholl analysis, a maximum projection of the z-stacks was computed, and binary masks were derived by manually tracing the dendrites. Using the SNT toolbox for analysis of neuronal morphology in ImageJ, skeletonized cell masks were analyzed at every 1.2 µm. Furthermore, soma size, primary dendrites and the total number of branch points were counted manually. For dendritic spine density analysis, high magnification images were obtained from dendritic segments with 0.14 µm pixel resolution with 0.14 µm z-steps. Dendritic spine density was counted on five dendritic segments (28-32 µm medium length) of each cell at an average distance of 66.2-70.0 µm from the soma.

### Behavioral procedures

#### Safety/Threat conditioning

Mice were handled for 10 minutes each day for three days before safety, threat or control conditioning. Subsequent conditioning took place over 3 days and recall took place 24 hours after the last conditioning session. Conditioning and recall sessions took place in behavioral chambers (MED Associates, Vermont, USA) housed within a sound-proof box and were recorded with infrared camera. All tone and shock deliveries were triggered via an Arduino and photometry and camera were triggered simultaneously via a Master-9 (A.M.P.I, Israel). Tones produced by the Arduino were in the form of buzzer-like square waves.

*Safety conditioning* consisted of a 2-minute habituation period followed by a delivery of 4 tone CS’s (continuous, 1000 Hz, 70 dB spl, 20 seconds + green LED) then 4 shock foot shock (2 seconds, 0.6 mA)). The CS and foot shock presentations were separated by a pseudo-random interval of (70-120 seconds). The order CS and foot shock presentations were reversed on day 2 (4 shocks followed by 4 tones at a pseudo-random interval). *Threat conditioning* consisted of the same habituation period and then 4 CS’s co-terminating with a foot shock which was delivered in the same pseudo-random interval. *Tone alone controls* received 4 CS presentations at the same intervals as safety conditioned mice but did not receive any foot shocks. The recall session for safety conditioned mice and tone alone controls took place in the original conditioning context (metal bar floors with no added odor) and consisted of 10 presentations of CS in the absence of foot shock at 1-minute intervals. The retrieval session for threat conditioned mice took place in a new context (plastic floor with peppermint odor) and consisted of the same 10 CS presentations in the absence of foot shock. Freezing during the 20 second CS presentation and the 20 second period prior to CS presentation (preCS) was measured using ANY-maze video tracking system (version 4.99z). ANY-maze “freeze score” was re-termed for this paper to “movement score” since greater freeze score values corresponded to greater movement. This re-terming was done to avoid potential misinterpretation of this value. The criterion used to define freezing onset was a movement score below a threshold 60 for at least 2 seconds and terminated when movement score rose above a threshold of 70. The freezing threshold movement score of 60 was subtracted from the movement traces in Figure S4I-X so that movement score values below zero represent freezing time points while values above zero represent non-freezing time points.

#### Optofluid drug infusion procedure

D1:cre mice were injected with GCaMP6f in the TS and optofluidic fibers were implanted 0.1mm above injection site. After 4-6 weeks of viral expression mice were safety conditioned for 3 days. Then, on day 4 and 6 mice were infused with 0.2 µl of saline or TTX (5 ng/ml in saline) respectively at a rate of 0.1 µl per minute 15 minutes prior to drug training session. Recall session consisted of four 2 second 0.6 mA shocks at 1-minute intervals followed by ten 20 second safety cues (continuous, 1000 Hz, 70 dB, 20 seconds + green LED) delivered at 1-minute intervals. Calcium traces for safety cue and shock in all photometry experiments, including optofluidic drug infusions, were averaged and waveform analysis was used to determine times when the 95% confidence interval did not contain a z-score of 0 and where the averaged traces were significantly different from each other (*85*). This approach provides within-subject analysis of behavior and photometry results and decreases the variability caused by differences in learning or GCamP6f expression between mice.

#### Open Field

Mice were allowed to explore a 40cm x 40cm square arena while being recorded from above for 10 min. Videos were then processed in ANY-maze video tracking software (version 4.99z) to determine distance traveled and mean speed.

#### Shock or tone calcium response assessment under anesthesia

The following day after conditioning recall, mice expressing GCaMP6f and optical fibers implanted in the TS were anesthetized with 4% isoflurane. Mice were then placed on a stereotaxic frame and anesthesia was maintained at 2% isoflurane. For shock delivery, mouse hind paws were lightly clamped with smooth alligator clips and attached to a Coulbourne shock generator. Five shocks (1 sec, 0.6 mA) were delivered at 1-minute intervals. For tone delivery, a speaker was positioned near the mouse’s head and five tones (continuous square wave 1000 Hz, 70 dB SPL, 20 sec) were played at 1-minute intervals. GCaMP responses were recorded and averaged together. After completion of the experiment, mice were euthanized for histological validation.

### Photometry data acquisition

A commercial fiber photometry apparatus (Doric Lenses) was used for recording calcium signals. The system consisted of a console connected to a computer, photometric multipliers tubes (PMTs), a four-channel programmable LED driver, and two LEDs at 405 and 465 nm connected to fluorescence dichroic mini cubes via 0.22NA 200 µm optical fibers. We measured bulk fluorescence from the TS using a single optical fiber for both delivery of excitation light and collection of emitted fluorescence. To measure GCaMP6f fluorescent changes reflecting neural activity we used 465 nm excitation. Meanwhile 405nm excitation was used to measure changes reflecting artifacts such as autofluorescence, movement, and bleaching. Files were analyzed using MATLAB (MathWorks).

### Quantification and statistical analysis

#### Statistical methods

Data were plotted with Prism 9 (GraphPad) or MATLAB and analyzed statistically with Prism 9 (GraphPad). Parametric tests (t tests; ANOVAs followed by Tukey, Dunnett’s or Holm-Sidak post hoc tests) were used in cases where data met assumptions of normality and equal variance. In cases where data did not conform to these assumptions, non-parametric tests (Wilcoxon signed-rank or Mann-Whitney rank sum tests; Friedman’s repeated-measures ANOVA on ranks followed by Student-Newman-Keuls post hoc tests) were used. The violin plots are displayed as median ± quartiles and line graphs are expressed as mean ± standard error of the mean (SEM). Data in the text are expressed as mean ± SEM. Detailed statistical analysis can be seen is Supplementary Table 2.

#### Analysis and quantification of photometry data

To facilitate the comparison of fiber photometry traces across multiple animals, we applied a multi-step process. First, the raw 465 nm and 405 nm channel data were individually converted into normalized change of fluorescence (ΔF/F) traces using the formula:

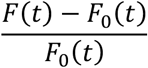

where *f*(*t*) is the original signal and *f*_0_(*t*) is a control trace generated by fitting the signal using least mean squares method. The ΔF/F trace from the 405 nm channel was then subtracted from the ΔF/F trace from the 465 nm channel to correct for artifacts of movement, bleaching, and autofluorescence. Next, standardized z-scores were calculated using the expression:

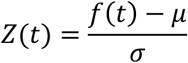

where μ is the mean and σ is the standard deviation of the ΔF/F trace. The z-scores were normalized around a specific time window. For CS delivery, the selected time window began 20 seconds prior to CS onset and ended 20 seconds after CS offset. For foot shock and threat recall termination, the selected time window begins 5 sec prior to event onset and ended 10 seconds after shock onset. The average z-score for the preCS or pre-event period was then subtracted from all points within the time window and the resultant traces were then averaged together across animals. AUCs for each trace were computed by numerically integrating the curve using the trapezoidal method. To determine if any observed sex differences in the statistical tests were of sufficient power, a standard threshold of 0.8 was used.

To determine time periods of significance during CS or shock presentation, bootstrapped confidence intervals (bCI) were computed for each trace based on a previously published method(*1*), and bars were plotted above traces to illustrate periods where z-scores were or significantly different between traces or significantly different from 0 and either above 0.5 or below -0.5. Briefly, waveforms were randomly resampled with replacement (1000 iterations) to obtain bootstrapped means. 95% confidence interval limits were derived based on the percentiles of the sampled distribution and expanded by a factor of 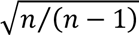 to counter small sample narrowness bias. Periods where confidence interval limits did not contain 0 (baseline) for at least 1/3 s were considered significant transients(*85*).

## Supporting information

All Supplementary Figures, Tables, and Movies

## Author Contributions

Conceptualization: A.S., D.S., and M.C.M

Methodology: A.S., D.S., M.C.M., C.L., S.J.C., S.C.

Formal analysis: A.S., S.C., and N.S.

Experiments and data collection: A.S., S.J.C., E.M., and A.F.

Writing - original draft: A.S., and S.J.C.

Writing - review & editing: A.S., S.J.C., D.S., M.C.M., C.L., N.S., S.C., E.M., and A.F.

Supervision: D.S. and M.C.M.

We thank Rui Costa and Eleanor Simpson for comments and suggestions.

Competing interest: Authors declare that they have no competing interests.

## Data and Materials Availability

### Lead contact

Further information and requests for resources and reagents should be directed to and will be fulfilled by the lead contact, David Sulzer.

### Materials availability

This study did not generate new unique reagents.

### Data and code availability

The source code for all photometry analysis is posted on GitHub: https://github.com/DSulzerLab/Doric_Photometry_Matlab_Analysis

All the data used in this study are available for download at: https://doi.org/10.6084/m9.figshare.26224232

