## Supplementary material for "Safety learning induces postsynaptic potentiation of direct pathway spiny projection neurons in the tail of the striatum": All Supplementary Figures, Tables, and Movies

Science Advances
AAAS

### Supplementary Materials for

#### Safety learning induces postsynaptic potentiation of direct pathway spiny projection neurons in the tail of the striatum

Adrien Stanley *et al.*

**This PDF file includes:**

Figs. S1 to S5
Movies S1 to S2

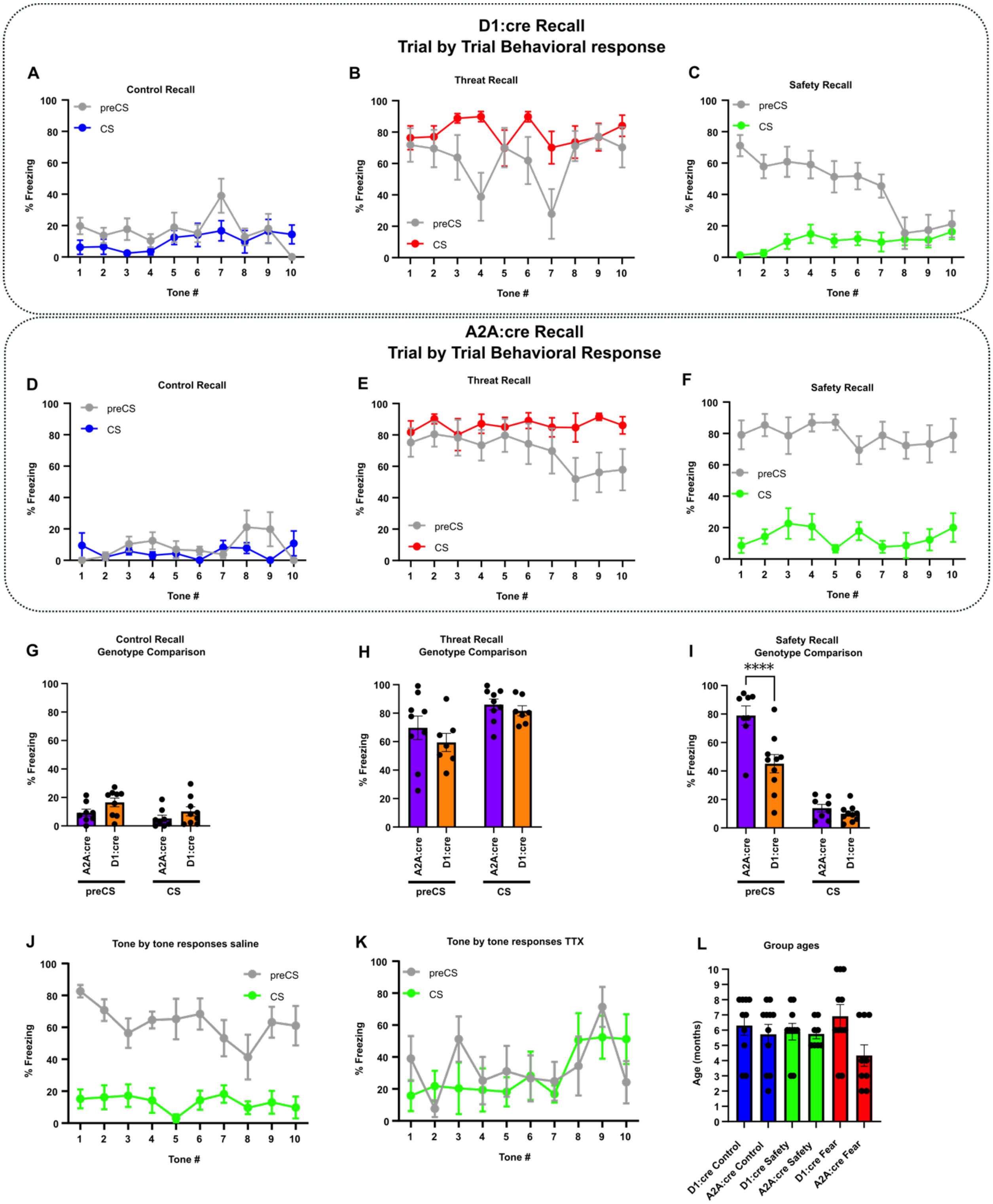


**Figure S1. Behavioral response comparisons between D1:cre and A2A:cre mice following sound-associative learning**. **(A-C)** Average percent time freezing behavior trial by trial during recall in **(A)** control conditioned, **(B)** threat conditioned and **(C)** safety conditioned D1:cre mice. **(D-F)** Average percent time freezing behavior trial by trial during recall in **(D)** control conditioned, **(E)** threat conditioned and **(F)** safety conditioned A2A:cre mice. **(G-I)** Comparison of mean freezing between D1:cre and A2A:cre genotypes for **(G)** control conditioned (Two-way ANOVA followed by Šídák's Test: genotype factor (F(1, 15)=2.863 P=0.1113), CS phase factor (F(1, 15)=8.299 P=0.0114), interaction factor (F(1, 15)=0.3998 P=0.5367), D1:cre, n=9; A2A:cre n=8), **(H)** threat conditioned (Two-way ANOVA followed by Šídák's Test: genotype factor (F(1, 14)=1.032 P=0.3268), CS phase factor (F(1, 14)=16.59 P=0.0011), interaction factor (F(1, 14)=0.3859 P=0.5445), D1:cre, n=7; A2A:cre n=9), and **(I)** safety conditioned mice (Two-way ANOVA followed by Šídák's Test: genotype factor (F(1, 16)=11.38 P=0.0039), CS phase factor (F(1, 16)=145.9 P<0.0001), interaction factor (F(1, 16)=12.99 P=0.0024), D1:cre, n=10; A2A:cre n=8). **(J-K)** Trial by trial behavioral response during recall of **(J)** vehicle infused mice (Two-way ANOVA followed by Šídák's Test: Tone number factor (F(2.776,11.10)=1.455 P=0.2792), CS phase factor (F(1, 4)=75.05 P=0.0010), interaction factor (F(3.023,12.09)=1.709 P=0.2176), n=5). **(K)** TTX infused mice (Two-way ANOVA followed by Šídák's Test: Tone number factor (F(3.108,12.43)=1.911 P=0.1789), CS phase factor (F(1, 4)=0.3341 P=0.5942), interaction factor (F(2.746,10.98)=1.471 P=0.2756), n=5). **(L)** Age of all mice for each genotype and group (one-way ANOVA: F(5, 53=1.737) P=0.1423, Control D1:cre n=10; Control A2A:cre n=11; Threat D1:cre n=11; Threat A2A:cre n=9; Safety D1:cre n=10; Safety A2A:cre n=8). All data was represented as mean ± SEM; ****p < 0.0001.

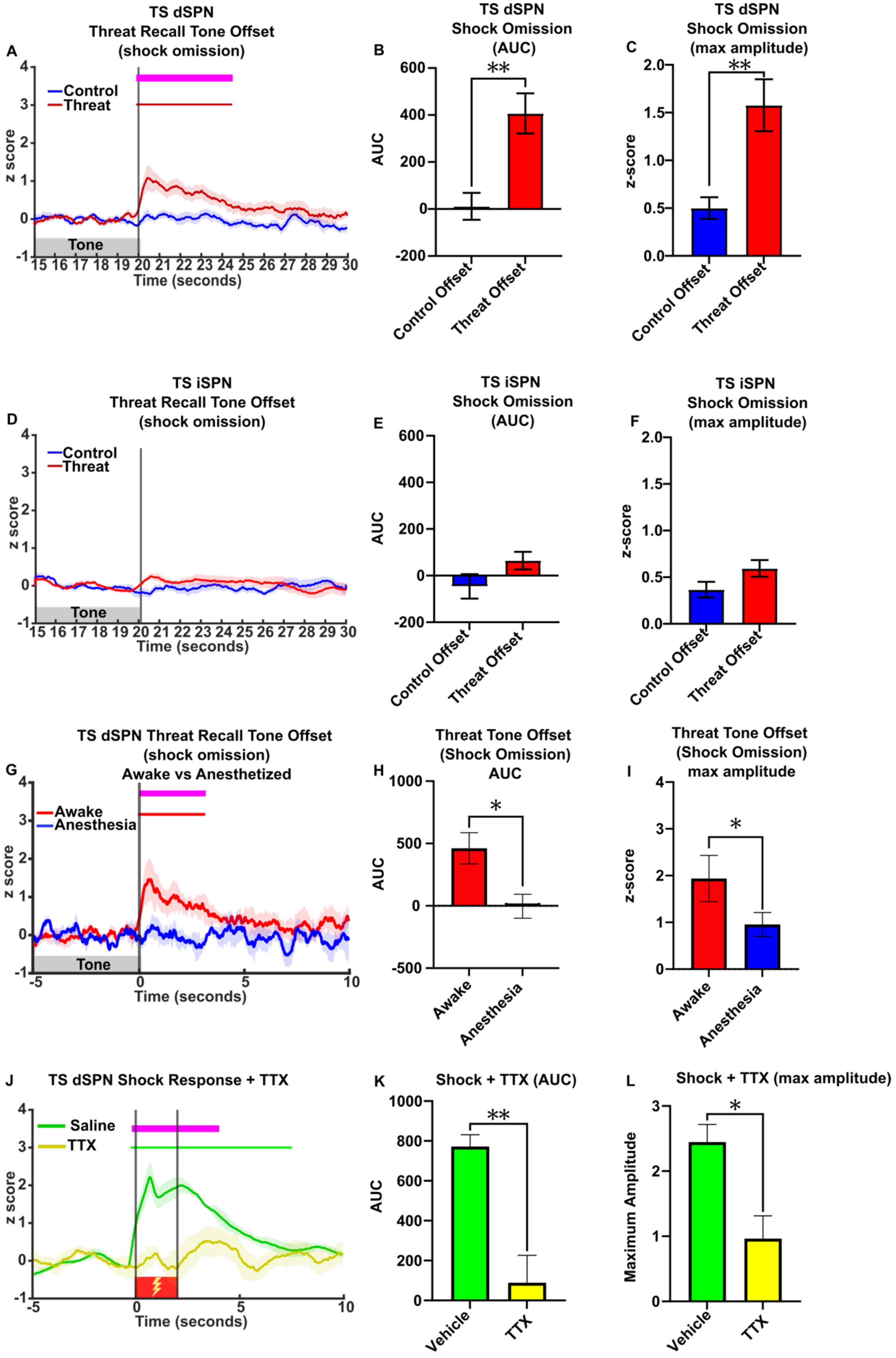
A
TS dSPN
Threat Recall Tone Offset
(shock omission)
Control
Threat
z score
Tone
Time (seconds)
B
TS dSPN
Shock Omission
(AUC)
AUC
**
Control Offset
Threat Offset
C
TS dSPN
Shock Omission
(max amplitude)
z-score
**
Control Offset
Threat Offset
D
TS iSPN
Threat Recall Tone Offset
(shock omission)
Control
Threat
z score
Tone
Time (seconds)
E
TS iSPN
Shock Omission
(AUC)
AUC
Control Offset
Threat Offset
F
TS iSPN
Shock Omission
(max amplitude)
z-score
Control Offset
Threat Offset
G
TS dSPN Threat Recall Tone Offset
(shock omission)
Awake vs Anesthetized
Awake
Anesthesia
z score
Tone
Time (seconds)
H
Threat Tone Offset
(Shock Omission)
AUC
AUC
*
Awake
Anesthesia
I
Threat Tone Offset
(Shock Omission)
max amplitude
z-score
*
Awake
Anesthesia
J
TS dSPN Shock Response + TTX
Saline
TTX
z score
Time (seconds)
K
Shock + TTX (AUC)
AUC
**
Vehicle
TTX
L
Shock + TTX (max amplitude)
Maximum Amplitude
*
Vehicle
TTX

**Figure S2. TS SPN responses to shock and shock omission**. Related to Figure 3. **(A)** TS dSPN average neural response to recall threat tone termination (red) and control tone termination (blue). Vertical line indicates tone offset (n=9). Bars above transients show periods significantly different from z-score of 0 for each condition ($p < 0.05$), and magenta bar shows periods of significance between conditions. **(B)** TS dSPN average AUC of recall threat tone offset and tone-alone control offset. (Unpaired t-test, p=0014; control, n=9; threat, n=9). **(C)** TS dSPN average maximum amplitude of recall threat tone offset and tone-alone control offset (Unpaired t-test, p=0.0021; control, n=9; threat, n=9). **(D)** TS iSPN average neural response to recall threat tone termination (red) and control tone termination (blue). Vertical line indicates tone offset (n=9). **(E)** TS iSPN average AUC of recall threat tone offset and tone-alone control offset. (Unpaired t-test, p=0.1098; control, n=7; threat, n=8). **(F)** TS iSPN average maximum amplitude of recall threat tone offset and tone-alone control offset. (Unpaired t-test, p=0.0787; control, n=7; threat, n=8). **(G)** Neural response to threat recall tone termination in awake (grey) and anesthetized (blue) mice (n=5). Bars above transients show periods significantly different from z-score of 0 for each condition ($p < 0.05$), and magenta bar shows periods of significance between conditions. **(H)** TS dSPN average AUC for 5 seconds following threat tone offset (paired t-test, p=0.018; n=5). **(I)** TS dSPN average maximum amplitude for 5 seconds following threat tone offset (paired t-test p=0.042; n=5). **(J)** Unpaired shock induced TS dSPN calcium response following vehicle and TTX administrations. Bars above transients show periods significantly different from z-score of 0 for each condition ($p < 0.05$), and magenta bar shows periods of significance between conditions (n=5). **(K)** TTX abolished TS dSPN AUC during shock (0-5 seconds from shock onset; paired t-test, p=0.005). **(L)** TTX abolished TS dSPN maximum amplitude during shock (0-5 second window from shock onset; paired t-test, p=0.033). All data was represented as mean ± SEM; *$p < 0.05$, **$p < 0.01$, ***$p < 0.001$.

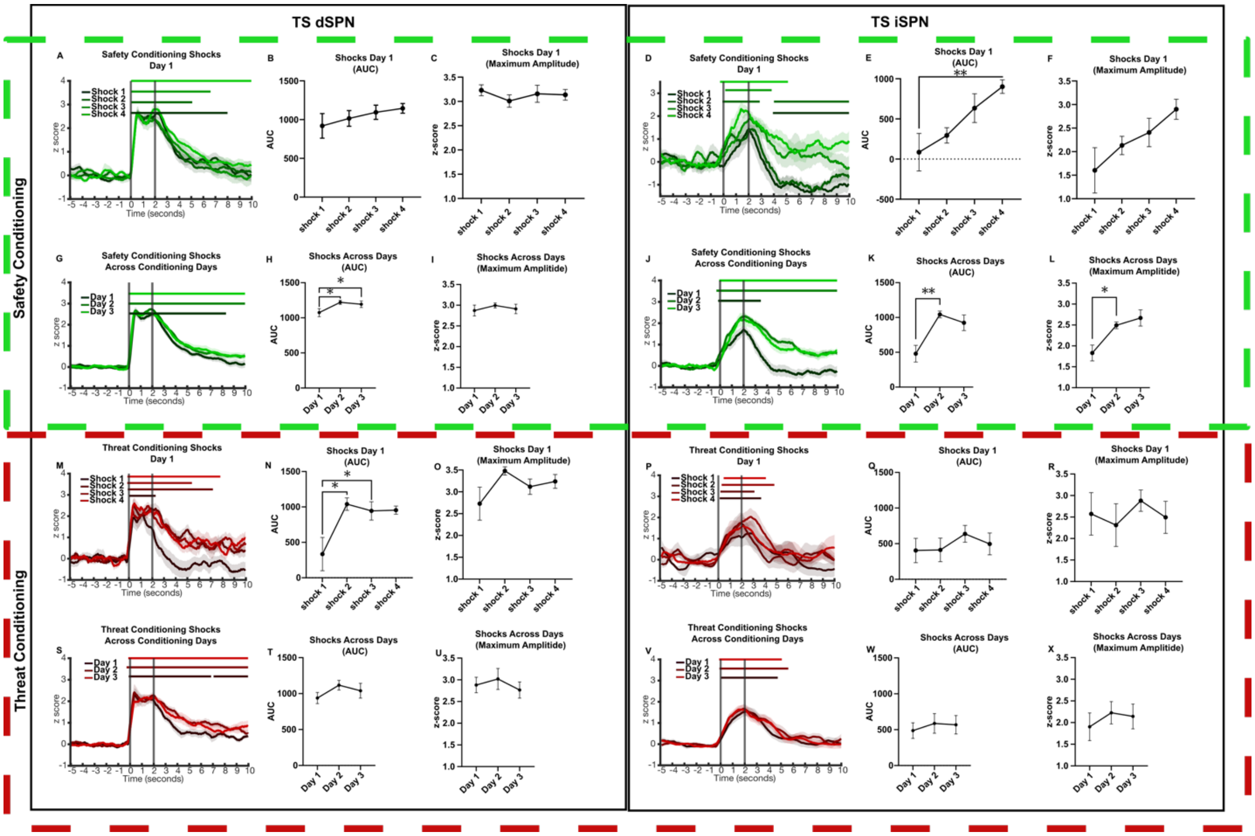


**Figure S3. TS SPN shock-related plasticity**. Related to Figure 2 and 3**. (A)** TS dSPN calcium response to unpaired shocks during the first day of safety conditioning. Average TS dSPN calcium response to day 1 unpaired shocks quantified in **(B)** AUC (one-way Anova with pairing, F (2.126, 19.14) = 1.112, P=0.3526, n=10) and **(C)** maximum amplitude (one-way Anova with pairing, F (2.320, 16.24) = 3.331, P=0.0555, n=10). **(D)** TS dSPN calcium response to unpaired shocks during the first day of safety conditioning. Average TS dSPN calcium response to day 1 unpaired shocks quantified in **(E)** AUC (one-way Anova with pairing followed by Dunnets test, F (1.895, 13.26) = 8.773, P=0.0041, n=8) and **(F)** maximum amplitude (one-way Anova with pairing, F (2.273, 18.18) = 0.5634, P=0.6002, n=8). **(G)** TS dSPN calcium response to unpaired shocks across days of safety conditioning. Average TS dSPN calcium response to unpaired shocks across days quantified in **(H)** AUC (one-way Anova with pairing followed by Dunnets test, F (1.226, 8.584) = 8.543, P=0.0148, n=10) and **(I)** maximum amplitude (one-way Anova with pairing followed by Dunnets test, F (1.179, 8.255) = 6.452, P=0.0304, n=10). **(J)** TS iSPN calcium response to unpaired shocks across days of safety conditioning. Average TS iSPN calcium response to unpaired shocks across days quantified in **(K)** AUC (one-way Anova with pairing followed by Dunnets test, F (1.512, 13.61) = 8.659, P=0.0060, n=8) and **(L)** maximum amplitude (one-way Anova with pairing followed by Dunnets test, F (1.618, 14.57) = 1.039, P=0.3633, n=8). **(M)** TS dSPN calcium response to unpaired shocks during the first day of safety conditioning. Average TS dSPN calcium response to day 1 unpaired shocks quantified in **(N)** AUC (one-way Anova with pairing followed by Dunnets test, F (1.762, 14.09) = 6.802, P=0.0103, n=10) and **(O)** maximum amplitude (one-way Anova with pairing, F (1.594, 12.75) = 2.588, P=0.1214, n=10). **(P)** TS dSPN calcium response to paired shocks during the first day of safety conditioning. Average TS dSPN calcium response to day 1 paired shocks quantified in **(Q)** AUC (one-way Anova with pairing, F (2.181, 17.45) = 0.7515, P=0.4971, n=9) and **(R)** maximum amplitude (one-way Anova with pairing, F (2.410, 19.28) = 0.4896, P=0.6543, n=9). **(S)** TS dSPN calcium response to paired shocks across days of safety conditioning. Average TS dSPN calcium response to paired shocks across days quantified in **(T)** AUC (one-way Anova with pairing, F (1.679, 13.43) = 2.002, P=0.1768, n=9) and **(U)** maximum amplitude (one-way Anova with pairing, F (1.393, 11.15) = 1.136, P=0.3330, n=9). **(V)** TS iSPN calcium response to paired shocks across days of safety conditioning. Average TS iSPN calcium response to paired shocks across days quantified in **(W)** AUC (one-way Anova with pairing, F (1.806, 14.44) = 0.4551, P=0.6238, n=9) and **(X)** maximum amplitude (one-way Anova with pairing, F (1.852, 14.82) = 0.7903, P=0.4629, n=9). All data was represented as mean ± SEM; *p < 0.05, **p < 0.01. Bars above transients show periods significantly different from z-score of 0 for each condition (p < 0.05).

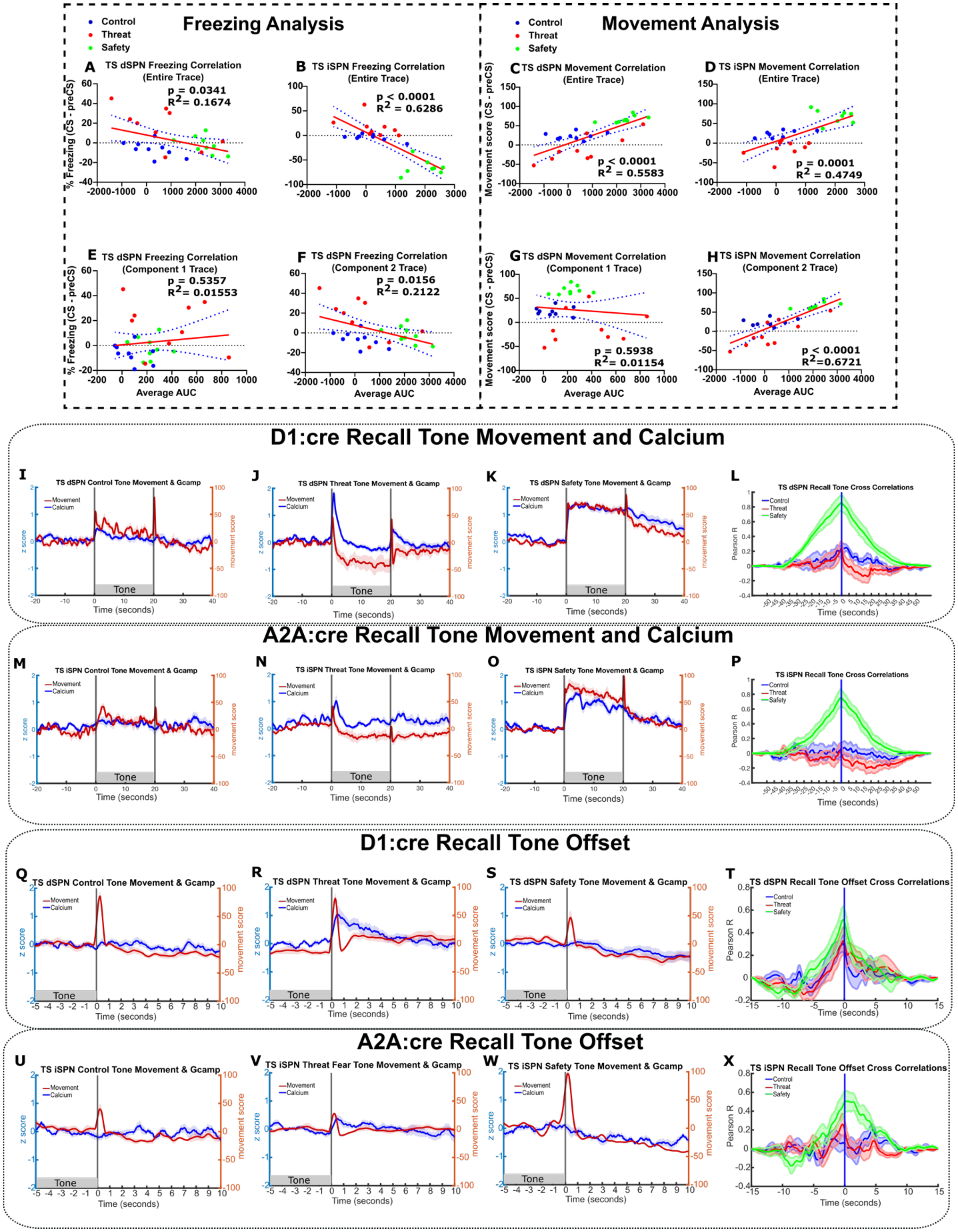


**Figure S4. TS SPN activity partially correlates with locomotor activity**. Related to Figure 3**.** **(A)** TS dSPN relationship between neural activity during entire CS period and freezing difference (CS - preCS). A linear regression revealed a significant negative relationship ($R^2$ = 0.1674, p =

0.0341). **(B)** TS iSPN relationship between neural activity during entire CS period and freezing difference (CS - preCS). A linear regression revealed a significant negative correlation ($R^2$ = 0.6286, p < 0.0001). **(C)** TS dSPN relationship between neural activity during entire CS period (0 – 20 seconds) and movement score difference (CS - preCS). A linear regression revealed a significant positive correlation ($R^2$ = 0.5583, p < 0.0001). **(D)** TS iSPN relationship between neural activity during entire of CS period (0 – 20 seconds) and movement score difference (CS - preCS). A linear regression revealed a significant positive relationship ($R^2$ = 0.4749, p < 0.0001). **(E)** TS dSPN relationship between neural activity during onset of CS period (0 – 2 seconds) and freezing difference. A linear regression revealed no correlation ($R^2$ = 0.01553, p = 0.5357). **(F)** TS dSPN relationship between neural activity during later CS period (2 – 20 seconds) and freezing difference (CS - preCS). A linear regression revealed a significant negative relationship ($R^2$ = 0.2122, p = 0.0156). **(G)** TS dSPN relationship between neural activity during onset CS period (0 – 2 seconds) and movement score difference (CS - preCS). A linear regression revealed no significant correlation ($R^2$ = 0.1154, p = 0.5938). **(H)** TS dSPN relationship between neural activity during later CS period (2 – 20 seconds) and movement score difference (CS - preCS). A linear regression revealed a significant positive correlation ($R^2$ = 0.6721, p < 0.0001). **(I-K)** Overlapped traces representing calcium activity (blue trace) and movement score (red trace) during recall for D1:cre mice that were **(I)** control conditioned, **(J)** threat conditioned, **(K)** and safety conditioned. X-axis represents time from tone onset and vertical lines represent tone onset and offset. **(L)** Cross correlation between movement and calcium activity during recall in D1:cre mice for control (blue), threat (red), and safety (green). **(M-O)** Overlapped traces representing calcium activity (blue trace) and movement score (red trace) during recall for A2A:cre mice that were **(M)** control conditioned, **(N)** threat conditioned, **(O)** and safety conditioned. Vertical lines represent tone onset and offset. X-axis represents time from tone onset. **(P)** Cross correlation between movement and calcium activity during recall in A2A:cre mice for control (blue), threat (red), and safety (green). X-axis represents time from tone onset and vertical lines represent tone onset and offset. **(Q-S)** Overlapped traces representing calcium activity (blue trace) and movement score (red trace) during tone offset for D1:cre mice that were **(Q)** control conditioned, **(R)** threat conditioned, **(S)** and safety conditioned. X-axis and vertical line represent time from tone offset. **(T)** Cross correlation between movement and calcium activity during recall tone offset in D1:cre mice for control (blue), threat (red), and safety (green). **(U-W)** Overlapped traces representing calcium activity (blue trace) and movement score (red trace) during tone offset for A2A:cre mice that were **(U)** control conditioned, **(V)** threat conditioned, **(W)** and safety conditioned. X-axis and vertical line represent time from tone offset. **(X)** Cross correlation between movement and calcium activity during recall tone offset in A2A:cre mice for control (blue), threat (red), and safety (green).

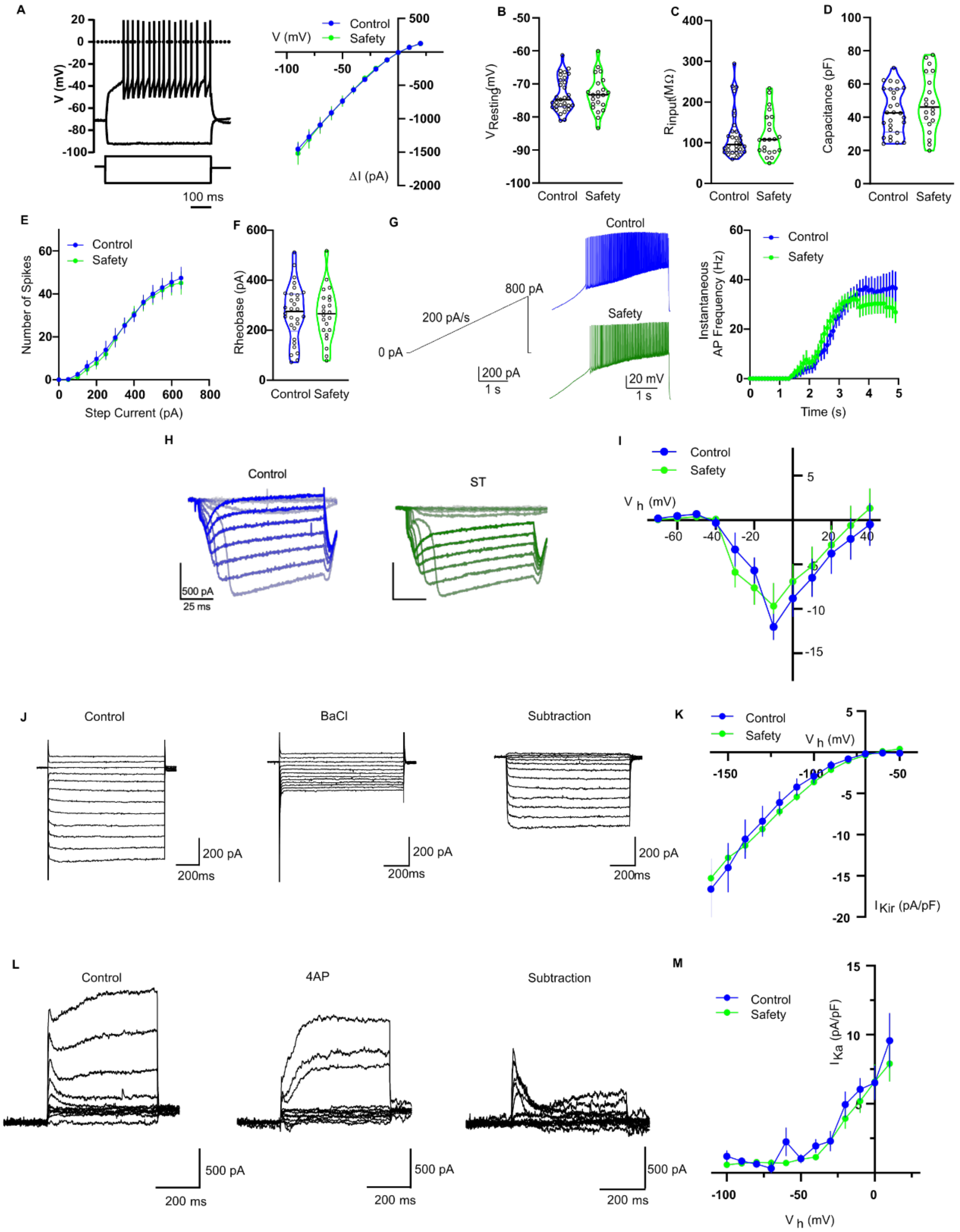

**Figure S5. Electrophysiological responses of TS dSPNs**. Related to Figure 5**. (A)** Left: Representative voltage responses in TS dSPNs evoked by 500 ms current injection. Right: Voltage vs. current relationship for safety conditioned and control TS dSPN (Control: N=7, n=24, Safety: N=7, n=19). **(B)** Resting membrane potential for safety conditioned and control TS dSPN (Control: N=7, n=24, Safety: N=7, n=19). **(C)** Input resistance for safety conditioned and control TS dSPNS (Control: 141.6 ± 4.35 MΩ, Safety: 109.6 ± 10.49 MΩ). **(D)** Capacitance for safety conditioned and control TS dSPNS (Control: N=12, n=37, 55.0 ± 4.24 pF, Safety: N=12, n=36, 68.06 ± 4.64 pF). **(E)** Number of spikes evoked by stepwise increases in current (-10 pA to +390 pA in 20 pA increments) (Control: N=10 n=21, Safety: N=7 n=16). **(F)** Rheobase for safety conditioned and control TS dSPNS (Control: 213.6 ± 16.18 pA, Safety: 287.2 ± 23.07 pA). **(G)** Left: Representative traces in response to a train of action potentials delivered by ramp current injection (4 s duration, 0-800 pA at 200 pA/s). Right: Action potential generation at different time points during ramp current injection (Two-way ANOVA followed by Bonferroni's test, time factor ($F(49, 2100) = 9.632$ $p<0.0001$), group factor ($F(1, 2100) = 128$ $p<0.0001$), interaction factor ($F(49, 2100) = 1.71$, $p=0.0017$), Control: N=7, n=26, Safety: N=7, n=18). **(H)** Representative traces of total $Ca^{2+}$ currents evoked by incremental voltage steps applied to TS D1 SPNs from control (left) or safety conditioned (right) mice. **(I)** Current-voltage curves of $Ca^{2+}$ currents between control and safety groups (Two-way ANOVA followed by Bonferroni's test, voltage factor ($F(11, 252) = 9.634$ $p<0.0001$), group factor ($F(1, 252) = 0.3486$, $p=0.5554$), interaction factor ($F(11, 252) = 0.3778$, $p=0.9638$), Control: N=5. n=10; Safety: N=5, n=14). **(J)** Example of inward rectifying potassium (Kir) currents. Currents were isolated by 500 µM $BaCl_2$. **(K)** Average current vs voltage relationship of $BaCl_2$-sensitive Kir currents between control and safety conditioned group (Two-way ANOVA followed by Bonferroni's test, voltage factor ($F(11, 228) = 39.82$ $p<0.0001$), group factor ($F(1, 228) = 0.1619$ $p=0.6878$), interaction factor ($F(11, 228) = 0.2328$, $p=0.9951$), Control: N=3 n=10, Safety: N=3 n=11). **(L)** Example of A type potassium ($K_a$) currents. Currents were isolated by 2 mM 4-aminopyridine (4AP). **(M)** Average current vs voltage relationship of $K_a$ currents (Two-way ANOVA followed by Bonferroni's test, voltage factor ($F(10, 176) = 25.76$ $p<0.0001$), group factor ($F(1, 176) = 2.162$, $p=0.6878$), interaction factor ($F(10, 176) = 0.4075$, $p=0.9418$), Control: N=3 n=8, Safety: N=3 n=10). The violin plots are displayed as median ± quartiles. All other data was represented as mean ± SEM.

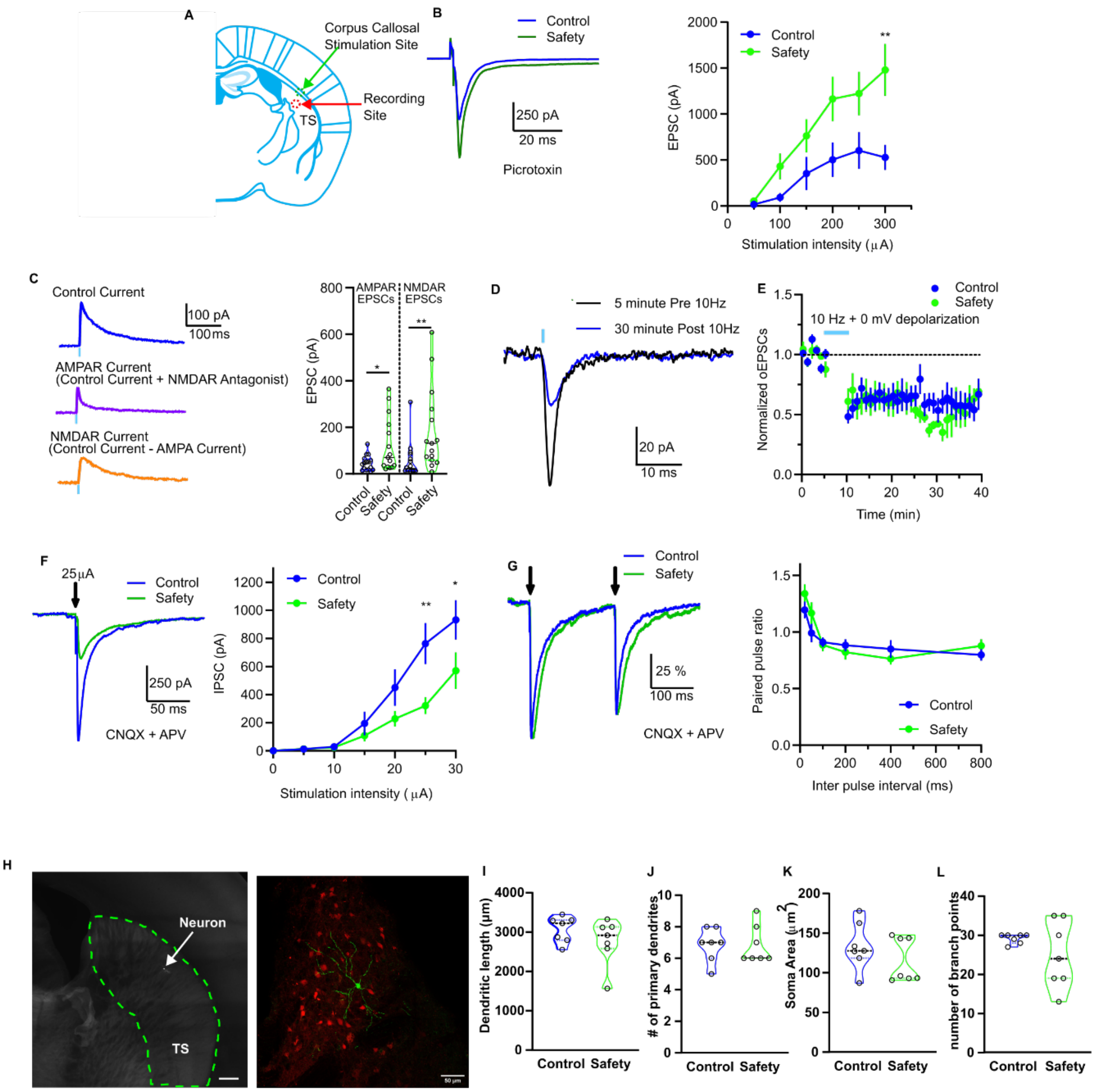


**Figure S6. Electrophysiological responses of TS dSPNs**. Related to Figure 5**. (A)** Schematic illustrating corpus callosal stimulation. **(B)** Left: Representative traces of evoked EPSCs from TS dSPNs by corpus callosal stimulation in safety conditioned and control mice. Right: Average TS dSPN EPSCs elicited by corpus callosal stimulation (Two-way ANOVA followed by Bonferroni's test, current factor ($F(5, 131) = 9.461$ $p<0.0001$), group factor ($F(1, 131) = 23.99$ $p<0.0001$), interaction factor ($F(5, 131) = 1.543$, $p=0.1809$), Control: N=3, n=12, Safety: N=3, n=12). **(C)** Left: Representative traces demonstrating the pharmacological isolation of NMDAR components at +60 mV by bath perfusing 25 µM APV. Right: Percent NMDAR contribution to optically evoked response (Mann Whitney test for AMPAR, $p<0.05$; Mann Whitney test for NMDAR, $p<0.01$, Control: N=7, n=14, Safety: N=6, n=15). **(D)** Sample optically evoked thalamo-striatal LTD in TS dSPN in control mice. **(E)** Time-course plot displaying long-term depression induced by 10 Hz thalamic stimulation paired with 0 mV depolarization. (two-way ANOVA, time factor ($F(36, 420) = 6.069$ $p<0.0001$), group factor ($F(1, 420) = 1.931$ $p=0.1654$), interaction factor ($F(36, 420) = 0.6331$, $p=0.9529$), Control: N=4, n=8, Safety: N=3, n=8). **(F)** Left: Representative traces of evoked IPSCs at 25 µA from control and safety trained groups. Right: Average TS dSPN IPSCs elicited by local stimulation (two-way ANOVA followed by Bonferroni's test, current factor ($F_{(6, 200)} = 26.03$ $p<0.0001$), group factor ($F_{(1, 200)} = 13.21$ $p=0.0004$), factor interaction factor ($F_{(6, 200)} = 2.550$ $p=0.0211$), Control: N=6, n=16, Safety: N=5, n=15). **(G)** Left: Representative traces of normalized paired-pulse evoked IPSCs from control and safety trained groups. Right: Paired-pulse ratio between control and safety trained group (Two-way ANOVA, ratio factor ($F_{(5, 222)} = 14.00$ $p<0.0001$), group factor ($F_{(1, 222)} = 0.9250$ $p=0.337$, interaction factor ($F_{(5, 222)} = 1.319$ $p=0.257$), Control: N=5, n=16, Safety: N=5, n=15). **(H)** Left: Representative location of patched TS dSPN filled with biocytin. Scale bar 300 µm. Right: higher magnification view of biocytin filled cell (green) and tdTomato expressing TS dSPNs in red. **(I-L)** Scholl analysis of **(I)** dendritic length (Mann Whitney test, p=0.3829, Control: N=4, n=7, Safety: N=6, n=8), **(J)** number of primary dendrites (Mann Whitney test, p=0.8427, Control: N=4, n=7, Safety: N=6, n=8), **(K)** soma surface area, (Mann Whitney test, p=0.535, Control: N=4, n=7, Safety: N=6, n=8), **(L)** and number of branch points (Mann Whitney test, P=0.535, Control: N=4, n=7, Safety: N=6, n=8). Violin plots in are displayed as median ± quartiles. All other data was represented as mean ± SEM ***p < 0.001.

#### Table S1. Statistical analysis of sex differences.

| Relevant figure | measurement | genotype | group | phase | male mean | female mean | male N | female N | p value | power | required N for power of 0.8 |
|---|---|---|---|---|---|---|---|---|---|---|---|
| 2C | Freezing | D1 | tone | preCS | 17.16 | 15.725 | 5 | 4 | 0.83127022 | 0.05464554 | 656 |
| | | | | CS | 7.64 | 13.475 | 5 | 4 | 0.40363107 | 0.16096361 | 30 |
| 2G | Freezing | D1 | safety | preCS | 39.2 | 49.08333333 | 4 | 6 | 0.48126404 | 0.17302693 | 30 |
| | | | | CS | 6.475 | 12.3 | 4 | 6 | 0.1142868 | 0.77150484 | 6 |
| 2K | Freezing | D1 | Fear | preCS | 50.2666675 | 73.375 | 4 | 4 | 0.04214153 | 0.69526381 | 5 |
| | | | | CS | 82.88889 | 76.025 | 4 | 4 | 0.40539373 | 0.09693682 | 58 |
| 2E | Freezing | A2A | tone | preCS | 9.314285714 | 9.4 | 7 | 1 | 0.99188227 | 0.05000934 | 122127 |
| | | | | CS | 4.471428571 | 11.1 | 7 | 1 | 0.37653241 | 0.12791064 | 17 |
| 2I | Freezing | A2A | safety | preCS | 83.76666667 | 64.55 | 6 | 2 | 0.24394419 | 0.53655126 | 6 |
| | | | | CS | 14.91666667 | 10.85 | 6 | 2 | 0.55004783 | 0.08537815 | 58 |
| 2M | Freezing | A2A | Fear | preCS | 65.15 | 71 | 2 | 7 | 0.78947021 | 0.90381991 | 4 |
| | | | | CS | 78 | 88.35714286 | 2 | 7 | 0.29562115 | 0.54404232 | 6 |
| 2N | % freezing diffrence | D1 | Tone only | | 23.185 | 23.37 | 4 | 5 | 0.97859225 | 0.05032384 | 9370 |
| | | | Safety | | -32.725 | -36.78333333 | 4 | 6 | 0.75636501 | 0.0683476 | 188 |
| | | | Fear | | 32.6222225 | 2.65 | 4 | 4 | 0.02807618 | 0.91875588 | 4 |
| 2P | % freezing difference | A2A | Tone only | | -4.842857143 | 1.7 | 7 | 1 | 0.39102954 | 0.12290194 | 18 |
| | | | Safety | | -68.85 | -53.7 | 6 | 2 | 0.27707457 | 0.33436887 | 9 |
| | | | Fear | | 12.85 | 17.35714286 | 2 | 7 | 0.79407842 | 0.1029914 | 42 |
| 3N, 3O, S4E | AUC Component 1 | D1 | safety | | 278.9671261 | 229.2587808 | 3 | 6 | 0.60492582 | 0.09184329 | 68 |
| | AUC Component 1 | D1 | fear | | 275.421314 | 344.3220376 | 5 | 5 | 0.75930694 | 0.05900064 | 396 |
| | AUC Component 1 | D1 | tone only | | 135.0663656 | 24.12130989 | 6 | 4 | 0.14038112 | 0.25182505 | 19 |
| S4F | AUC Component 2 | D1 | safety | | 2809.856321 | 1909.008581 | 3 | 6 | 0.02103637 | 0.87759815 | 4 |
| | AUC Component 2 | D1 | fear | | -337.8404735 | -307.3378423 | 5 | 5 | 0.97168485 | 0.05018419 | 19177 |
| | AUC Component 2 | D1 | tone only | | 473.8970308 | 86.36729578 | 6 | 4 | 0.48307584 | 0.09395823 | 80 |
| 4N, 4O, S4G | AUC Component 1 | A2A | safety | | 161.6486823 | 281.747332 | 6 | 2 | 0.14918469 | 0.30706554 | 9 |
| | AUC Component 1 | A2A | fear | | 94.59134101 | 100.6259176 | 2 | 7 | 0.96181692 | 0.0517747 | 1199 |
| | AUC Component 1 | A2A | tone only | | 65.57659202 | 11.97962645 | 7 | 2 | 0.5636468 | 0.07828933 | 78 |
| S4H | AUC Component 2 | A2A | safety | | 1812.974543 | 1782.343657 | 6 | 2 | 0.94519981 | 0.05044455 | 4403 |
| | AUC Component 2 | A2A | fear | | 13.67259917 | 118.4278654 | 2 | 7 | 0.85492357 | 0.10390326 | 42 |
| | AUC Component 2 | A2A | tone only | | 437.9511048 | 52.094779 | 7 | 2 | 0.522061 | 0.0852364 | 63 |

**Table S2. Statistics Table.** Spreadsheet containing the detailed statistics performed for every experiment in the manuscript.

Link to spreadsheet with Table S2

**Movie S1. Representative behavioral and TS D1 SPN calcium response to a safety associated tone at recall.** Video showing behavioral response at during a tone at safety recall during preCS (before LED activation) and CS (during LED activation). Safety cue onset and offset are indicated by vertical lines in the calcium trace.
Link to Supplementary Video 1

**Movie S2. Representative behavioral and TS D1 SPN calcium response to a threat associated tone at recall.** Video showing behavioral response at during a tone at threat recall during preCS (before LED activation) and CS (during LED activation). Threat cue onset and offset are indicated by vertical lines in the calcium trace.
Link to Supplementary Video 2
